# Feeding-dependent tentacle development in the sea anemone *Nematostella vectensis*

**DOI:** 10.1101/2020.03.12.985168

**Authors:** Aissam Ikmi, Petrus J. Steenbergen, Marie Anzo, Mason R. McMullen, Anniek Stokkermans, Lacey R. Ellington, Matthew C. Gibson

## Abstract

In cnidarians, axial patterning is not restricted to embryonic development but continues throughout a prolonged life history filled with unpredictable environmental changes. How this developmental capacity copes with fluctuations of food availability and whether it recapitulates embryonic mechanisms remain poorly understood. To address these questions, we utilize the tentacles of the sea anemone *Nematostella vectensis* as a novel paradigm for developmental patterning across distinct life history stages. As a result of embryonic development, *Nematostella* polyps feature four primary tentacles, while adults have 16 or more. By analyzing over 1000 growing polyps, we find that tentacle progression is remarkably stereotyped and occurs in a feeding-dependent manner. Mechanistically, we show that discrete *Fibroblast growth factor receptor b* (*Fgfrb*)-positive ring muscles prefigure the sites of new tentacles in unfed polyps. In response to feeding, a Target of Rapamycin (TOR)-dependent mechanism controls the expansion of *Fgfrb* expression in oral tissues which defines tentacle primordia. Using a combination of genetic, cellular and molecular approaches, we demonstrate that FGFRb regionally enhances TOR signaling activity and promotes polarized growth, a spatial pattern that is restricted to polyp but not to embryonic tentacle primordia. These findings reveal an unexpected plasticity of tentacle development, and show that the crosstalk between TOR-mediated nutrient signaling and FGFRb pathway couples post-embryonic body patterning with food availability.

## Introduction

Cnidarians such as sea anemones, corals, and hydrozoans have continuous developmental capacities^1–3^. Similar to plants, these early-branching animals can generate organs and body axes throughout their entire life. This developmental feature underlies diverse phenomena, such as secondary outgrowth formation^4^, asexual reproduction^5^, and branching in colonial species^6^. The ability to continuously build new body parts is comparable to regeneration, as both require activation of patterning mechanisms in a differentiated body plan. However, unlike regeneration induced by damage or injury, life-long organogenesis is subject to environmental modulation. This strategy allows cnidarians, like plants, to continuously adjust their developmental patterns to unpredictable fluctuations of food supply^2,7^. To determine how cnidarians deploy organogenesis across life history stages and whether this process recapitulates embryonic development or employs distinct regulatory mechanisms, we study post-embryonic tentacle development in the starlet sea anemone *Nematostella vectensis*.

Arrays of tentacles armed with stinging cells are a unifying feature of Cnidaria, with diverse species featuring distinct arrangements, morphologies, and numbers of tentacles^8–11^. In the typical cnidarian polyp *bauplan*, oral tentacles are simple extensions of the diploblastic body, forming appendages that feed, defend, and expand the surface area of the gastric cavity. Zoologists in the early 1900s described tentacle patterns in select species, some of which could exceed 700 tentacles (e.g. *Cereus pedunculatus*)^10^. The partial sequence by which tentacles are added over time was also reported, reflecting the existence of continuous axial patterning in polyps. In hydrozoans, this developmental property was primarily studied in the context of body plan maintenance and regeneration, where the oral tissue of *Hydra* features a Wnt/ß-catenin-dependent axial organizing capacity^12^. Still, how new morphological patterns are generated and how developmental patterning unfolds across distinct life history stages remain unknown. Interestingly, a link between tentacle morphogenesis and nutrition was recently observed in the sea anemone Aiptasia^13^, but the mechanistic basis of this food-nutrient dependency of tentacle development is still unsolved.

In the recent years, *Nematostella vectensis* has become an established cnidarian model in developmental biology due to its relative ease of laboratory spawning^14,15^, tractable developmental biology^16–20^, extensive regenerative capacity^21–24^, and robust molecular-genetic approaches^25–29^. *Nematostella* polyps can harbor a variable number of tentacles ranging from four to eighteen, but the common number in adulthood is sixteen^4,10^. During development, four tentacle buds simultaneously form in the swimming larvae and give rise to the initial appendages of the primary polyp^4^. The formation of tentacles involves coordination between both embryonic body layers. In developing larvae, an endodermal *Hox* code controls inner body segmentation, defining territories that are critical for tentacle patterning^20,30,31^. In the ectoderm, tentacle morphogenesis initiates from a thickened placode, followed by changes in epithelial cell shape and arrangement that drive tentacle elongation^4^. Nevertheless, we still have only a rudimentary understanding of the developmental relationship between embryonic and post-embryonic tentacles, or how tentacle progression integrates the nutritional status of the environment.

## Results

### Tentacle patterning in primary and adult polyps

*Nematostella* polyps possess two axes: one running from the pharynx to the foot (oral-aboral axis), and an orthogonal axis traversing the pharynx (directive axis; Figure 1A-D). Along the directive axis, the primary polyp displays mesenteries, internal anatomical structures that subdivide the body into eight recognizable radial segments (s1-s8) (Figure 1C and E; Figure S1)^20^. The four primary tentacles occupy stereotyped radial positions corresponding to segments s2, s4, s6 and s8 (n=131 polyps; Figure 1C and E). While this octo-radial body is maintained in adults, we found that the spatial pattern of tentacle addition generated three new features in mature polyps with 16-tentacles (Figure 1D and F; Figure S1). First, tentacles were arranged in a zigzag pattern forming two adjacent concentric crowns at the oral pole. Second, the boundaries between neighboring tentacles within the same segment were marked by the formation of short gastrodermal folds, enriched in F-Actin and called micromeres (Figure S1)^10^. Third, the arrangement of tentacles shifted from radial to bilateral organization with a defined number of tentacles in each segment. This included a single tentacle in the directive segments s1 and s5, two tentacles in segments s2, s8 s3 and s7, and three tentacles in segments s4 and s6 (n=108 polyps; Figure 1D and F). Together, these observations reveal that tentacle development in polyps derives from an intricate and reproducible spatial patterning system. In addition, we observed that adult polyps can grow more than 18-tentacles when they were not regularly spawned (Figure S2), suggesting a trade-off between resource allocation to reproduction and adult organogenesis.

**Figure 1:**
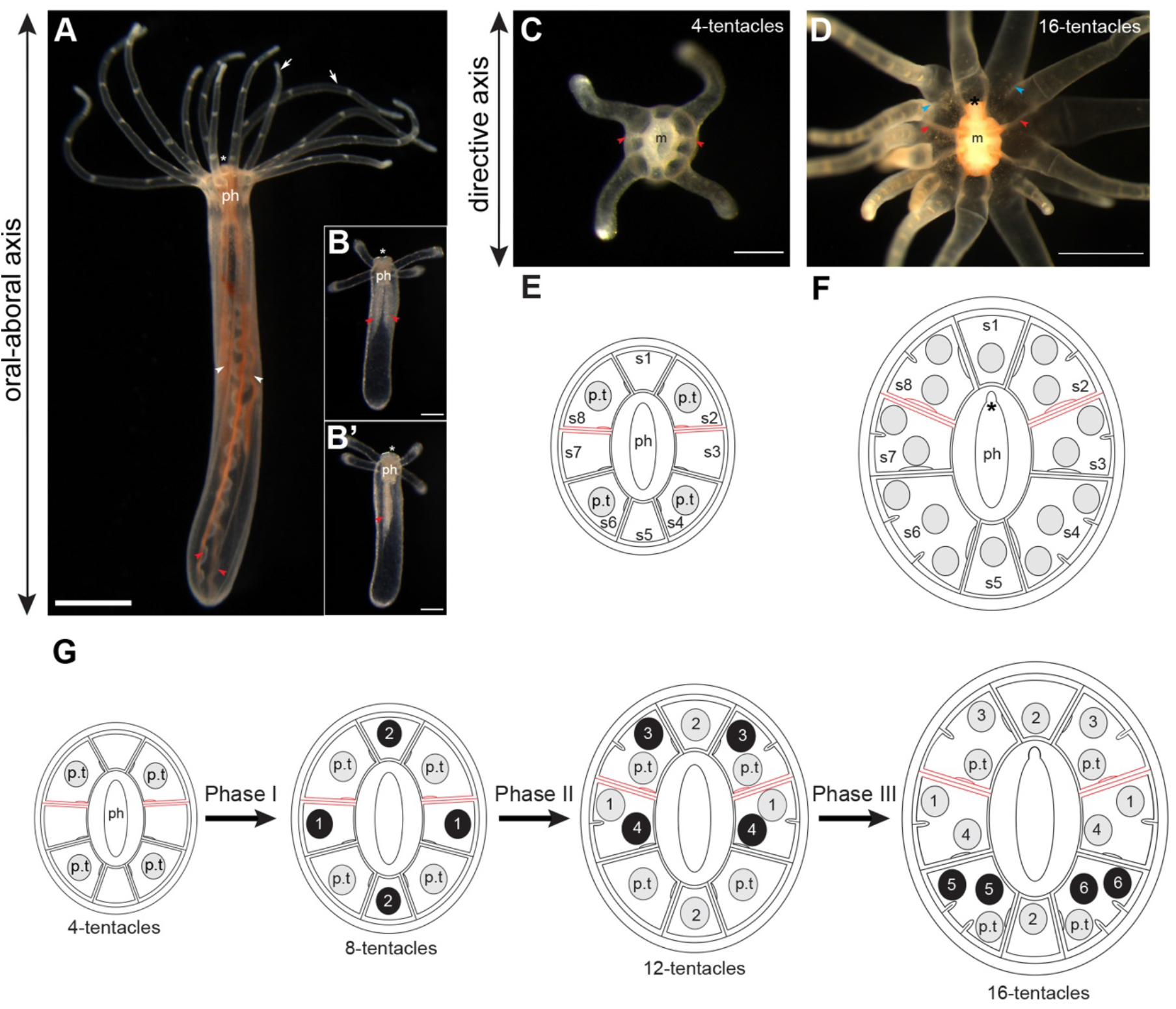
Body axes and tentacle arrangement in primary and adult polyps. **B**, Lateral views of an adult polyp bearing 12 tentacles and a four-tentacle primary polyp, respectively. White asterisk (*) indicates the position of the oral pole. The pharynx (*ph*) is attached to the body wall through eight endodermal mesenteries. In primary polyps, two primary mesenteries (*red arrowheads*) are well-developed along the oral-aboral axis and form a mirror image along the directive axis. In adults, these primary mesenteries maintain a growth advantage over the remaining mesenteries (*white arrowheads*). **B**’, 90° rotation of the primary polyp shown in **B.** Note that the position of primary mesenteries can serve as a landmark to orient the oral pole along the directive axis. White arrows show tentacles. Scale bars are 1mm and 100μm in **A** and **B/B’**, respectively. **C** and **D**, Oral views of a primary polyp and an adult with 16 tentacles, respectively. Both oral poles show eight segments and are oriented along the directive axis using the position of primary mesenteries (*red arrowheads*). Black asterisk (*) indicates the position of the siphonoglyph, a ciliated groove positioned at one end of the mouth (*m*). *Blue* arrowheads indicate two examples of boundaries between neighboring tentacles within segments. Scale bars are 250μm and 1mm in **C** and **D**, respectively. **E** and **F**, Diagrammatic cross-section through the oral pole, summarizing the arrangement of tentacles in primary polyps and 16 tentacles adults, respectively. The eight body segments are annotated from s1 to s8. Primary mesenteries are colored in red. Tentacles are depicted as grey discs. Primary tentacles (p.t) resulting from embryonic development are indicated. **G**, Summary of the representative pattern of tentacle addition showing the three phases. The sequence of budding events is indicated as black discs with a number.

### Feeding-dependent stereotyped tentacle formation

In the absence of nutrients, we found that *Nematostella* polyps arrested at the 4-tentacle stage and rarely developed additional tentacles (Figure S3). When food was available, however, primary polyps grew and sequentially initiated new tentacles in a nutrient-dependent manner, arresting at specific tentacle stages in response to food depletion (Figure S3). To build a spatio-temporal map of tentacle addition, we leveraged this feeding-dependent development to control the progression of tentacle stages. We examined 1102 fixed specimens collected between the 4- and 16-tentacle stages and recorded the placement and order of tentacle formation (Figure 2 and Figure 3). Despite using a non-isogenic strain of *Nematostella*, tentacle addition was surprisingly stereotyped and was dependent on two distinct budding modalities, which we term *cis*- and *trans*-budding. In both modes, tentacles were generated in pairs, either through simultaneous or consecutive budding events. In *trans*-budding, new tentacles formed in opposing segments along the directive axis (Figure 2A-C). In *cis*-budding, a pair of new tentacles developed in the same segment (Figure 3A and B). These budding patterns occurred in a distinct temporal sequence. Based on the spatio-temporal deployment of new tentacles, we conclude that the nutrient-dependent development of tentacles falls into three phases mediated by six pairs of budding events (Figure 1G).

**Figure 2:**
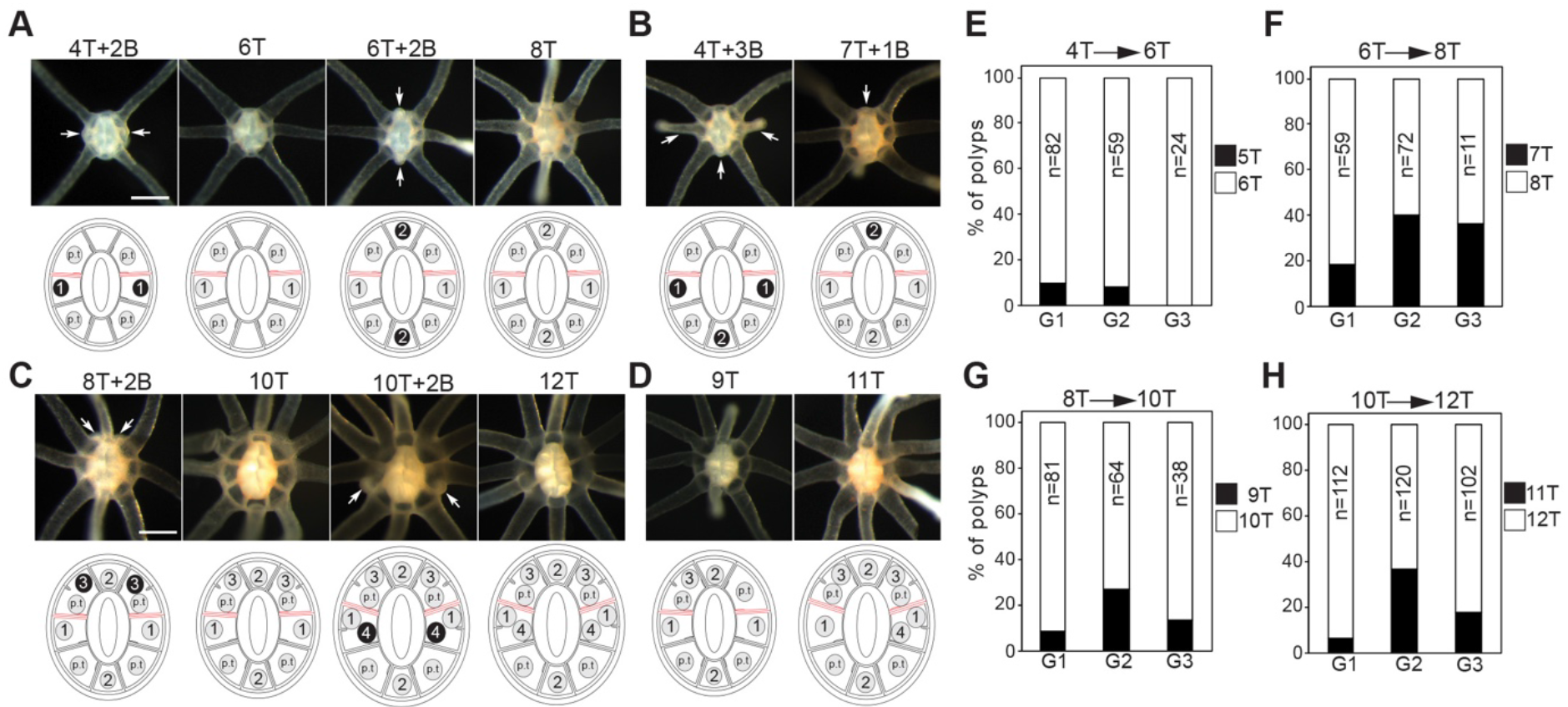
Phase I and II of tentacle addition. **A-D**, Oral views of fed polyps. **A** and **B**, Progression from 4 to 8 tentacles through dominant and alternative budding patterns, respectively. **C**, Progression from 8 to 12 tentacles. **D**, Polyps with 9 and 11 tentacles. *White* arrows show the sites of budding events. The number of tentacles (T) and buds (B) is indicated. Scale bars are 250μm. Diagram showing tentacle arrangement is provided under each image. Tentacles and buds are depicted as grey and black discs, respectively. The numbers inside the discs indicate the sequence of budding. The positions of primary tentacles (p.t) are shown. **E-H**, Quantification of tentacle number including buds in polyps from three independently growing groups (G1, G2 and G3). Tentacle progression is indicated on the top of each graph and the numbers of analyzed polyps are indicated. Source data are provided as a Source Data file.

**Figure 3:**
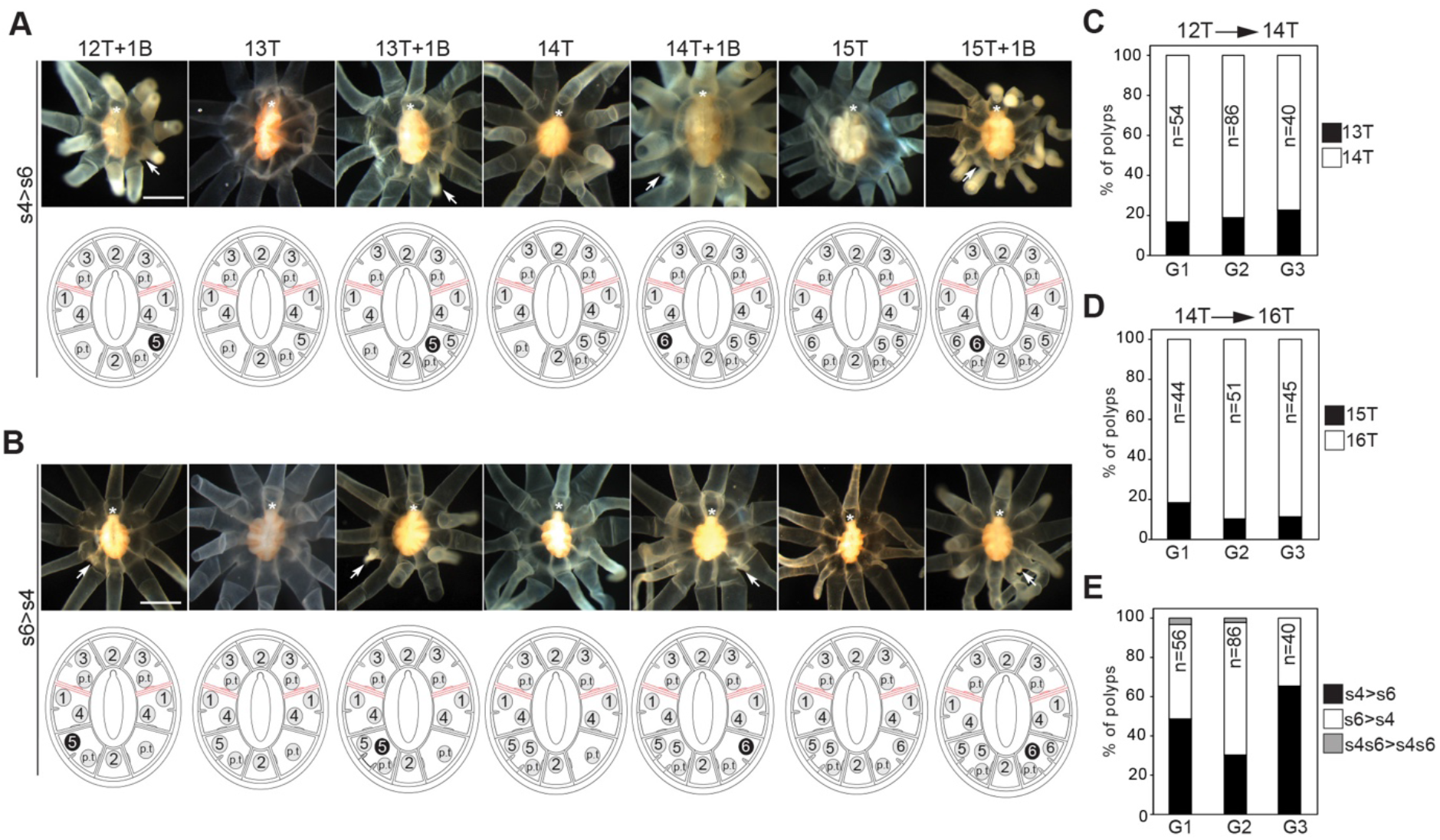
Phase III of tentacle addition. **A** and **B**, Oral views of fed polyps progressing from 12 to 16 tentacles. **A**, Cis-budding events taking place in s4 segment then s6 segment (s4>s6). **B**, Alternative pattern of cis-budding events (s6>s4). *White* arrows show the sites of budding events. The number of tentacles (T) and buds (B) is indicated. Scale bars are 500μm. Diagram showing tentacle arrangement is provided under each image. Tentacles and buds are depicted as grey and black discs, respectively. The numbers inside the discs indicate the sequence of budding. **C** and **D**, Quantification of tentacle number including buds in growing polyps. Tentacle progression is indicated on the top of each graph. **E**, Quantification of budding sequence in growing polyps. S4s6>s4s6 means two trans-budding events in s4 and s6 segments. Source data are provided as a Source Data file.

Phase I corresponded to the progression from the 4- to the 8-tentacle stage, where two *trans*-budding events took place in the tentacle-less segments s1, s3, s5 and s7 (Figure 2A-F). Tentacle buds initially developed in segments s3 and s7, followed by segments s1 and s5. The first *trans*-budding event was mostly simultaneous, while the second showed many cases of asynchrony (Figure 2B and F). At the end of phase I, each of the eight segments displayed a single tentacle. In phase II, a bilateral pattern of tentacles emerged as a result of two trans-budding events leading to 12-tentacle stage (Figure 2C-H). The third and fourth pairs of tentacles were added in segments s2/s8 and s3/s7, respectively. In contrast to phase I, trans-budding events were preceded by the formation of short gastrodermal folds (Figure S1). These new boundaries created territories for tentacle development within segments, with buds generated in stereotyped locations with respect to the pre-existing tentacles. During phase III, tentacle development relied on the *cis*-budding mode to proceed from the 12- to 16-tentacle stage (Figure 3A-D). The fifth and sixth pairs of tentacles were sequentially formed in segments s4 and s6, with no preferential bias in the order of deployment (Figure 3E).

Although the tentacle addition pattern was highly stereotyped, we also observed some temporal and spatial variability in tentacle pattern (Figure S4). A strong asynchrony in the development of tentacle pairs was the predominant variant observed during the progression from 4- to 12-tentacle stages (n=155/824). We also found animals where one sister pair of tentacles was skipped or formed in a neighboring segment. In other cases, the order of budding events was inverted or two pairs of bud formed simultaneously (n=66). During phase III, we found a few animals that used *trans*-budding instead of *cis*-budding (n=6/320). Collectively, these variants represented ~20.6% of all specimens scored, perhaps attributable to genetic variation, variation in nutrient uptake, and developmental plasticity in growing polyps.

### Feeding-induced tentacle development is size independent

To begin to elucidate the mechanisms that control tentacle addition, we focused on studying the transition from the 4- to the 6-tentacle stage. Following daily feedings of primary polyps, the first pair of buds appeared between day 4 and 6, depending on the amount of *Artemia* consumed (Figure S3). At the organismal scale, tentacle budding coincided with an approximate 2-fold increase in body size (Figure S5). To probe the relationship between body size and tentacle budding, we generated reduced-sized primary polyps by dividing four-cell stage embryos into two pairs of blastomeres and then allowing development to proceed (Figure 4A and B)^32^. Upon feeding, reduced-sized polyps developed tentacle buds at smaller oral circumferences compared to full-sized animals, indicating that the feeding-dependent tentacle development relies on a mechanism that scales in proportion to body size (Figure 4C-E).

**Figure 4:**
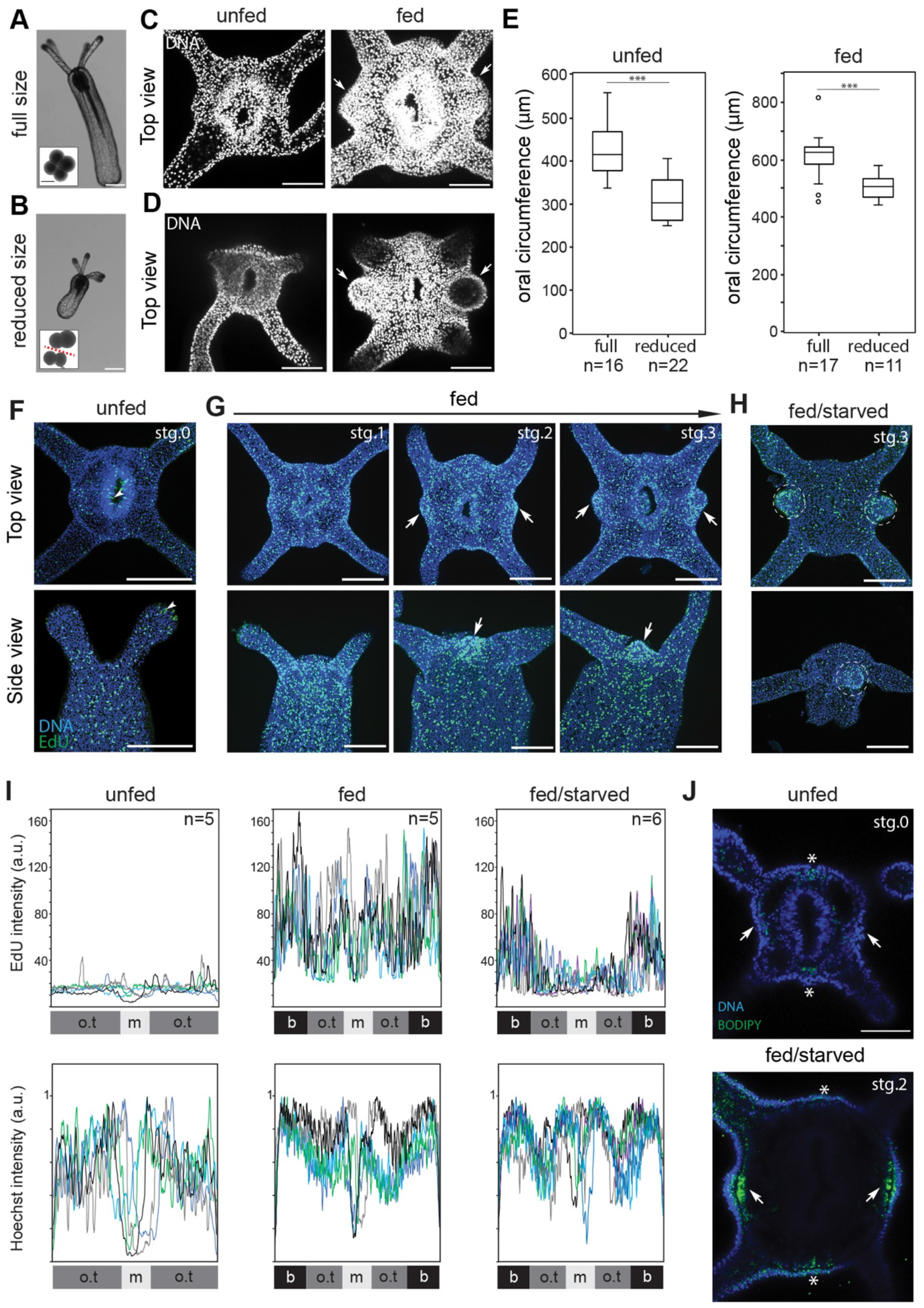
Polyp size and growth pattern during tentacle addition. **A**, Unfed primary polyp. Inset box shows a 4-cell stage embryo. **B**, Reduced-sized polyps resulting from blastomere isolation. Inset box shows 4-cell stage embryo divided into 2 pairs of blastomeres. Scale bars are 100μm. **C**, Confocal z-projection of the oral poles of full-sized polyps stained with Hoechst (white) in unfed and fed conditions. **D**, Confocal z-projection of the oral poles of reduced-sized polyps in unfed and fed conditions. White arrows indicate tentacle buds. Scale bars are 50μm. **E**, Quantification of oral circumferences of full sized, and reduced-sized polyps in unfed and fed conditions. Box plot values consist of the median (center line), upper and lower quartiles (upper and lower edges of box), and maximum and minimum values (whiskers). Outliers are represented in small circles. T-Student’s test *p*<0.0001. **F-H**, Confocal projections of animals stained for EdU incorporation (green) and with Hoechst (blue) to visualize S-phase cells and nuclei at the indicated feeding conditions. White arrows indicate localized enrichment of EdU incorporation. White arrowheads indicate examples of unspecific EdU-labelling at the tentacle tip and in the pharynx. Dashed circles show tentacle buds maintaining EdU while the animals were starved. Fed/starved means animals were fed for 3 days than starved for 4 days. Budding stages from 0 to 3 are indicated (see Figure S6). Scale bars are 100μm. **I**, Plots of EdU intensity and normalized Hoechst intensity across the oral tissue (o.t.) of segments s3/s7 and mouth (m) area in unfed, fed and fed/starved polyps. Bud (b) areas are also indicated. **J**, Confocal z-stacks of animals stained with BODIPY (green) and Hoechst (blue) at the indicated feeding conditions. White arrows and asterisk indicate the sites of the first and second trans-budding, respectively. Scale bar is 50μm. Source data are provided as a Source Data file.

To monitor growth during tentacle development, we labelled cells in S-phase with EdU incorporation and observed two distinct patterns of cell proliferation. Uniform S-phase labeling was induced in response to feeding. This was followed by a localized increase in cell proliferation in both cell layers of tentacle bud primordia (Figure 4F, G, and I; Figure S6). Bud-localized proliferation transformed the thin epithelial layers into a thickened outgrowth, generating the initial cellular organization associated with budding stages (Figure S6). During tentacle elongation, cell proliferation was uniform but excluded from tentacle tips, a region enriched with differentiating stinging cells^33,34^. These results show that tentacle morphogenesis in primary polyps is preceded by nutrient-dependent global growth. This is in turn followed by the formation of localized growth zones that mark the sites of the nascent buds.

To define the relationship between nutrient input and cell proliferation, we established a minimal feeding assay sufficient to induce tentacle budding. Under these conditions, polyps were fed for 3 days and then starved for 4 days until the first pair of buds developed. Interestingly, while we observed a dramatic reduction of uniform cell proliferation in the starved budded polyps, bud-localized cell proliferation was significantly less affected (Figure 4H and I). The different sensitivities to starvation suggest that distinct regulatory mechanisms control cell proliferation-associated with generalized organismal growth and region-specific tentacle budding. Interestingly, an enrichment of lipid droplets was detected in bud primordia, visualized with BODIPY and Oil Red O staining (Figure 4J; Figure S7). This localized energy storage could serve as a buffering mechanism to complete tentacle development under unpredictable fluctuations of food supply.

### TOR-dependent growth is required for tentacle formation

To test the role of growth in nutrient-dependent tentacle budding, we used Rapamycin to inhibit the TOR pathway, a growth-regulatory module that integrates multiple cellular inputs including nutrition, energy availability, and growth factor signaling^35^. Polyps were fed daily for 8 days while concomitantly treated with 1μM Rapamycin. As expected, control animals exhibited increased body size and developed tentacle buds (Figure S5). In contrast, animals treated with Rapamycin did not grow and failed to form new buds, although they internalized the orange pigment of *Artemia* as an indicator of successful feeding (Figure S5). Consistent with a growth defect, cell proliferation was dramatically reduced in Rapamycin treated animals (Figure 5C and D; Figure S8). Lack of bud formation was also observed when polyps were only treated with Rapamycin from day 3, during the expected period of tentacle budding (Figure S8). These results show a continuing requirement for TOR-dependent growth in post-embryonic tentacle development.

**Figure 5:**
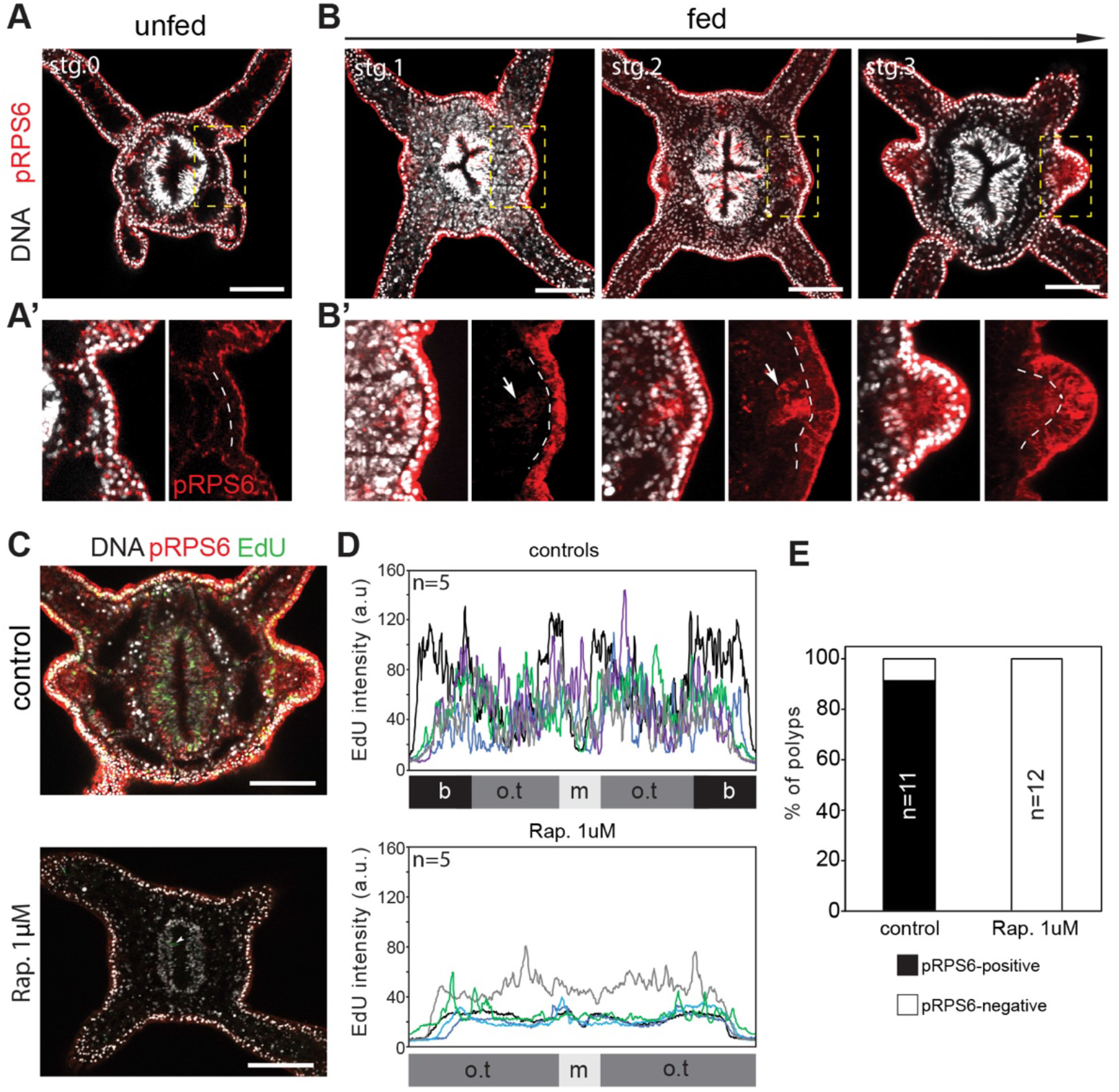
TOR pathway activity and function in fed polyps. **A**, Confocal z-section of the oral pole of unfed polyp stained with with an antibody against pRPS6 (red) and Hoechst to label nuclei (white). A’, Inset box is a zoom in of s3 segment **B**, Confocal sections of developing buds at sequential stages. B’, Inset boxes show a higher magnification of s3 segment for each stage. Dashed lines separate the outer- and inner-layer nuclei. Arrows indicate the enrichment of pRPS6 in the inner-layer of tentacle primordia. **C**, Confocal projection of the oral pole of 5 days fed control and 8 days fed Rapamycin-treated polyps stained for EdU incorporation (green), an antibody against pRPS6 (red) and with Hoechst (white). White arrowhead indicates an unspecific EdU-labelling in the pharynx **D**, Plots of EdU intensity in control and Rapamycin-treated polyps. **E**, Quantification of polyps showing PS6RP-postive tentacle primordia in control (n=11) and Rapamycin treated polyps (n=12). Scale bars are 50μm. Source data are provided as a Source Data file.

To visualize the activity of the TOR pathway, we utilized an antibody directed against a conserved phosphorylation motif in the 40S ribosome protein S6 (pRPS6; Figure S9)^36^. This protein is a direct substrate of ribosomal protein S6 kinase (S6K) and its phosphorylated form is a reliable marker for TOR Complex 1 and S6K activation^37^. In unfed polyps, cytoplasmic pRPS6 staining was not detected in either epidermal or gastrodermal layers, but pRPS6 immunoreactivity was mainly localized at the apical regions of epithelial cells (Figure 5A and A’). Following feeding, we observed ubiquitous cytoplasmic pRPS6 staining in the epidermal layer that reflected the active metabolic state of growing polyps (Figure 5B and B’; Figure S8). Interestingly, stronger cytoplasmic pRPS6 staining marked bud primordia in both germ layers and provided a clear molecular readout of tentacle patterning (Figure 5B and B’; Figure S8). Confirming the TOR dependence of RPS6 phosphorylation, fed animals exposed to rapamycin exhibited a dramatic reduction of pRPS6 and failed to phosphorylate RPS6 at presumptive tentacle primordia (Figure 5C and E; Figure S8). Taken together, these results show that feeding induces the organismal activation of the TOR pathway, which becomes spatially patterned and defines the location of tentacle primordia.

### *Fgfrb* expression marks tentacle primordia in polyps

Based on the results described above, we hypothesized that TOR-dependent organismal growth modulates the activity of developmental signaling pathways, which in turn generate a feedback loop that locally enhances TOR pathway activity and promotes polarized growth in tentacle primordia. FGFR signaling is an attractive candidate to mediate this function, as discrete cell clusters expressing *Fgfrb* prefigure the position of tentacle primordia in unfed polyps (Figure 6A and Movie S1)^38^. To define the identity of these orally-scattered cells, we generated a reporter line expressing eGFP under the control of ~8.7kb surrounding the promoter region of *Fgfrb* (Figure S10 and Movie S2). By combining fluorescent *in situ* hybridization of *Fgfrb mRNA* and α-eGFP immunostaining, we confirmed the overlap between the endogenous and transgenic *Fgfrb* expression in primary polyps (Figure 6C). Based on the morphology of eGFP-positive cells and F-actin staining, we assigned these *Fgfrb*-positive cell clusters to a sub-population of ring muscle cells, characterized by an epitheliomuscular architecture that display elongated basal processes associated with myofilaments (Figure 6C; Figure S10)^39^. Interestingly, these *Fgfrb*-positive cell clusters were established during larval development, indicating their pre-metamorphic origin (Figure S10). At the oral pole, the *Fgfrb-eGFP* line also labelled longitudinal muscles and pharyngeal cells, which is consistent with the endogenous expression pattern of *Fgfrb* (Figure S10).

**Figure 6:**
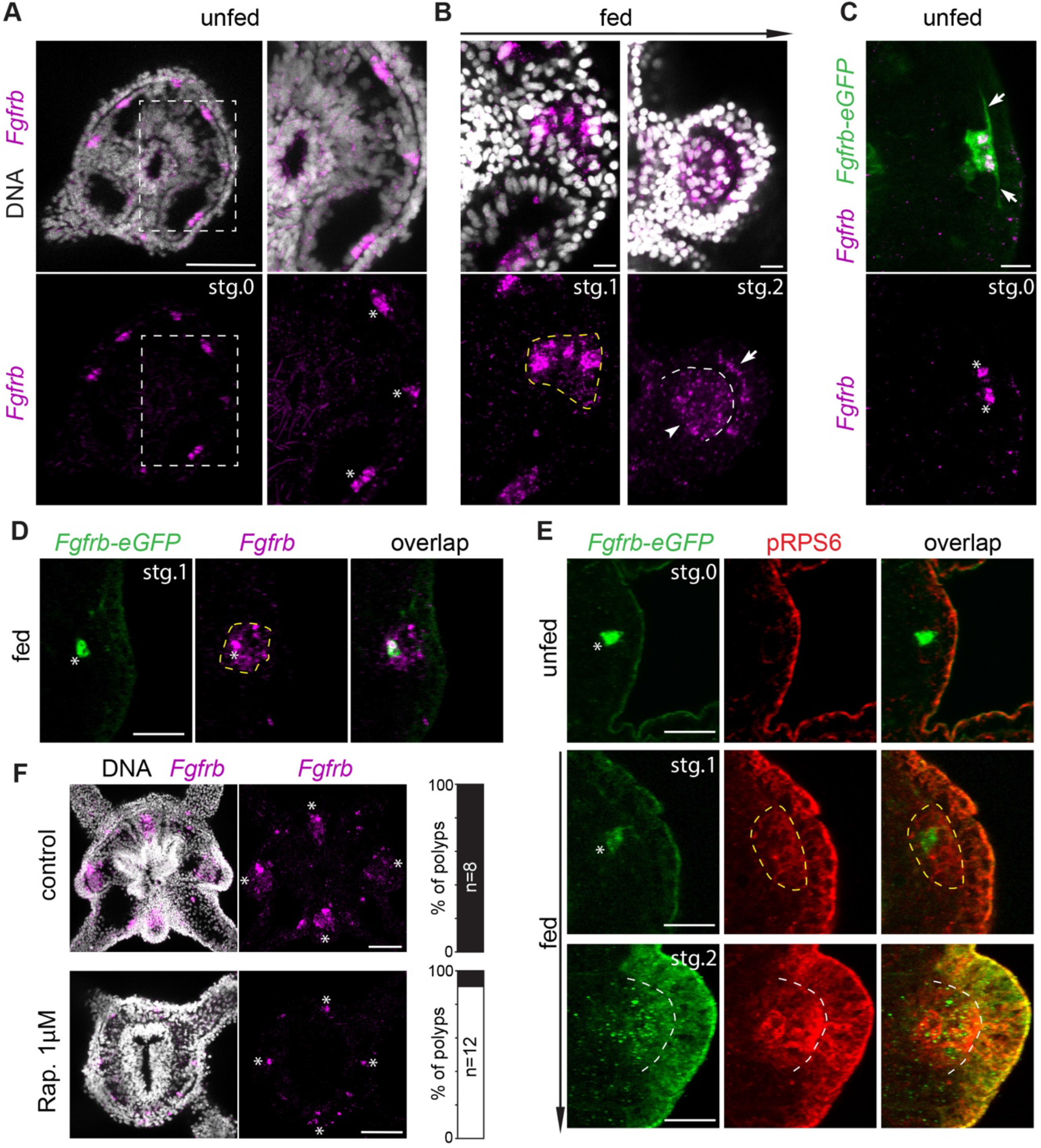
*Fgfrb*-positive ring muscles pre-mark the sites of tentacle primordia and TOR pathway is required for the expansion of *Fgfrb* expression during budding. **A** and **B**, Fluorescent *in situ* hybridization of *Fgfrb* in unfed and fed polyps. **A**, Confocal z-projections of the oral pole of unfed polyp showing *Fgfrb* expression (purple) and nuclei (white). Inset box is a zoom in of s3 segment. **B**, Confocal z-projections of developing buds at sequential stages. White dashed line separates the two body layers. White arrow and arrowhead indicate outer- and inner-body layer, respectively. **C and D**, Confocal z-projections of the *Fgfrb-eGFP* transgenic line stained with α-eGFP (green) and labeled for *Fgfrb* mRNA (purple) in unfed and fed polyp, respectively. White arrows show the elongated basal parts of ring muscles. White arrowheads indicate nuclear localization of *Fgfrb* mRNA in the eGFP-positive cells. Yellow dashed line delineates the expression domain of *Fgfrb* after feeding. E, Confocal z-projections of unfed and fed *Fgfrb-eGFP* polyps stained with α-eGFP (green) and pRPS6 antibody (red). Yellow dashed line delineates the domain of pRPS6-positive cells. F, Confocal projection of the oral pole of fed controls and fed Rapamycin-treated polyps stained with Hoechst (white) and labeled for *Fgfrb* mRNA (purple). Quantification is shown. The black bar indicates an expansion of *Fgfrb* expression in tentacle primordia while the white bar shows no change in *Fgfrb* expression. White Asterisks indicate *Fgfrb*-expressing cells in the oral segments. Scale bars are 100μm. Scale bars are: 50μm in A and F, 10μm in B and C, and 20μm in D and E. Source data are provided as a Source Data file.

To determine the effect of feeding on *Fgfrb* expression, we performed fluorescent *in situ* hybridization in growing and budding polyps (Figure 6A and B). In fed polyps, the expression of *Fgfrb* expanded from a small number of cells in the inner layer to a larger domain within the body segment (Figure 6A and B). This feeding-dependent expansion was nucleated around the initial *Fgfrb*-positive ring muscles as visualized by the different temporal dynamics between *Fgfrb* mRNA and *Fgfrb-eGFP* expression (Figure 6D). Interestingly, pRPS6-positive cells also adopted a similar organization surrounding the pre-feeding *Fgfrb*-positive cells (Figure 6E). During budding, both epidermal and gastrodermal cell layers showed an enrichment of *Fgfrb* expression that overlapped with pRPS6 staining in tentacle primordia (Figure 6B and E). Similar to TOR pathway activity, *Fgfrb* expression showed a segment-dependent regulation in response to feeding (Figure S10). Segment s3 and s7 were first to express this pattern, which mirrored the sequence of the first *trans*-budding event. Fed-polyps treated with 1μM Rapamycin did not show an expansion of *Fgfrb* expression while they maintained the discrete cell clusters expressing *Fgfrb* (Figure 6F). Taken together, these results show that there are at least two early events associated with the feeding-dependent tentacle development. First, ring muscle cells expressing *Fgfrb* pre-mark the sites of the morphogenetic changes in tentacle primordia. Second, TOR activity drives the feeding-dependent expansion of *Fgfrb* in neighboring cells.

### *Fgfrb* controls axial elongation and feeding-induced budding

To determine the role of *FGFR* signaling, we first treated fed-polyps with the FGFR signaling inhibitor SU5402 (Figure S11)^40^. While control and drug-treated polyps showed a similar size increase following 5 days of development, the expected spatial enrichment of pRPS6 staining and cell proliferation at tentacle primordia was not detected in SU5402-treated polyps (Figure S11). As SU5402 can inhibit both PDGF and VEGF signaling^41^, we next genetically disrupted *FGFRb* signaling. To do so, we generated independent mutant alleles of *Fgfrb* using the CRISPR/Cas9 system. Several alleles were isolated, including two putative null alleles *Fgfrb^mut1^* and *Fgfrb^mut2^*, which disrupted the first and second coding exons, respectively (Figure 7A). To minimize any possible CRISPR/Cas9 off-target effects that could be expressed in homozygous animals, we analyzed the phenotypes of F2 *trans*-heterozygous *Fgfrb^mut1^/Fgfrb^mut2^* individuals. Crossing F1 heterozygotes resulted in Mendelian ratios of viable F2 *trans*-heterozygous polyps, whose genotypes were confirmed by DNA sequencing (Figure S12). *Fgfrb^mut1^/Fgfrb^mut2^* animals exhibited a significantly reduced length of both body column and tentacles compared to sibling controls (Figure 7B and Figure S12). This phenotype most likely resulted from a failure of *Fgfrb* mutants to properly elongate during metamorphosis. In addition, *Fgfrb* mutants also displayed reduced septal filaments and defects in longitudinal tentacle muscle compared to their siblings (Figure S12). Nevertheless, these mutants exhibited the expected eight segments and four primary tentacles with cnidocyte enrichment at the tip, indicating normal proximo-distal patterning (Figure 7B and E).

**Figure 7:**
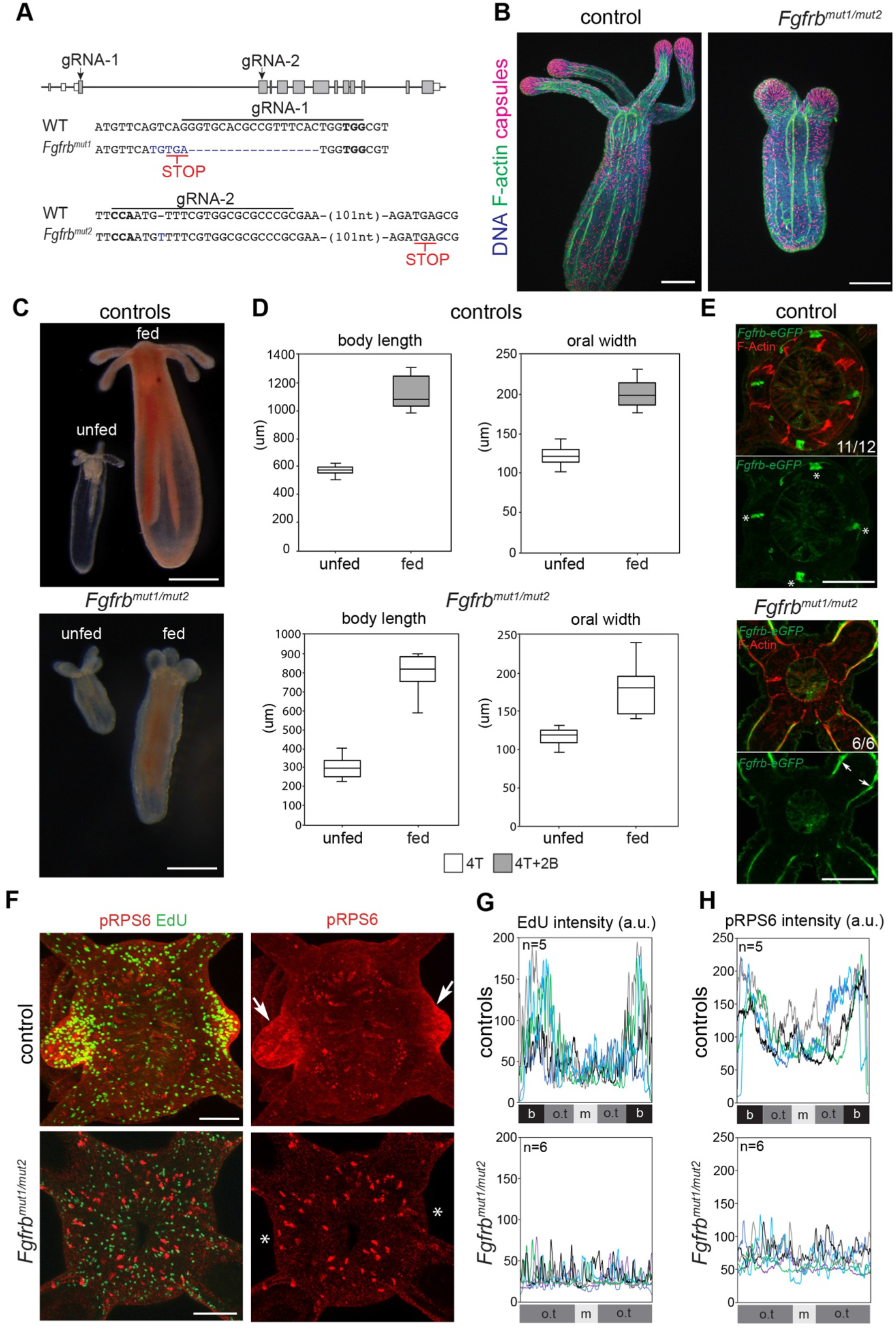
*Fgfrb* signaling is required for the feeding-dependent tentacle development. **A**, Gene model of the *Fgfrb* locus showing the target sites of gRNA1 and gRNA2 and resulting indel mutations that create premature stop codons (red). PAM motifs are in bold characters. **B**, Confocal z-projections of indicated polyps stained with Phalloidin (F-Actin, green) and DAPI (nuclei, blue). DAPI also stains mature nematocyst capsules (purple). Scale bars are 100μm. **C**, Image of unfed and fed indicated polyps. Scale bars are 250μm**. D**, Quantification of body length and oral width in control (unfed, n=15; fed, n=10) and *Fgfrb* mutant polyps (unfed, n=15; fed, n=8). The number of tentacles and buds is indicated. Box plot values consist of the median (center line), upper and lower quartiles (upper and lower edges of box), and maximum and minimum values (whiskers). **E**, Confocal z-projections of oral poles of indicated polyps stained with α-eGFP (green) and phalloidin (red). White asterisks indicate ring muscles expressing *Fgfrb*. Note that these discrete ring muscles cells, but not *Fgfrb*-positive longitudinal tentacle muscles, are missing in the mutant background. **F**, Confocal z-projections of oral poles of indicated polyps stained for EdU incorporation (green), an antibody against pRPS6 (red) and with Hoechst (blue). Scale bars are 50μm. White arrows indicate the first trans-budding event marked by pRPS6 in fed polyps. White asterisks (*) show the expected sites for tentacle bud formation. **G** and **H**, Quantification of EdU and pRPs6 intensity in control (Top) and mutant (bottom) polyps. In all panels, control animals are siblings of the *Fgfrb* mutants. Source data are provided as a Source Data file.

As *Fgfrb* mutants were viable, we tested their ability to resume development in response to feeding. Out of twenty hand-fed mutants, eight polyps consistently consumed food and exhibited growth, although a tripling of their initial body length took much longer (~six weeks versus two weeks in controls; Figure 7C-D). This delayed growth was not observed in SU5402-treated polyps (Figure S11), suggesting that this phenotype might be the result of disrupting *Fgfrb* function during embryonic-larval development. Still, despite the significant organismal growth of *Fgfrb* mutants, they failed to elongate the four primary tentacles and did not develop new tentacle buds (Figure 7C and D). Further, these mutants showed shorter and wider primary tentacles compared to controls. Consistent with a *Fgfrb*-dependent patterning defect in post-embryonic tentacle development, mutants lacked the oral *Fgfrb-positive* cell clusters as assessed with both endogenous mRNA and transgenic *Fgfrb* expression (Figure 7E and Figure S12). In addition, *Fgfrb* mutants did not exhibit pRPS6-positive domains or polarized growth in the expected segments (Figure 7F-H). Taken together, we conclude that *Fgfrb* signaling is dispensable for primary tentacle budding, but is essential to pattern TOR pathway activity and localized cell proliferation in subsequent tentacle primordia.

## Discussion

A diversity of organisms has evolved strategies to both sense and adapt to changes in their environment, resulting in the remarkable plasticity of development. Among animals, environmental plasticity has been widely described in ephemeral species (e.g. insects, worms and vertebrates). These animals typically experience specific periods in which development is responsive to environmental cues, producing long-lasting changes in either form or physiology^42^. In contrast, species with extreme longevity must continuously adjust their developmental behaviors to unpredictable fluctuations of food supply. This plasticity underlies diverse adaptive phenomena such as reversible body size changes in adult planarians^43^, or asexual reproduction in *Hydra*^44^. Along similar lines, here we uncovered the cellular and signalling mechanisms by which *Nematostella* polyps integrate the nutritional status of the environment to control post-embryonic tentacle development.

Our results establish the spatial map for tentacle addition in the sea anemone *Nematostella*. By examining a large number of polyps progressing from the 4- to the 16-tentacle stage, we showed that two budding modalities, *cis* and *trans*, drive tentacle addition in a stereotyped spatial pattern. While the mechanisms that direct the temporal sequence of budding are unknown, the initial step of this process is the formation of outgrowths from radial segments that lack primary tentacles. Once each segment developed a single tentacle, secondary tentacle territories sequentially emerged within select segments, except in the directive segments s1 and s5. These territories preceded tentacle development and were defined by the formation of short gastrodermal folds enriched in F-Actin. Based on these observations and as previously reported during Hox-dependent embryonic segmentation^20^, the formation of endodermal territories is a common theme in tentacle patterning. However, whether Hox genes play a central role in subdividing pre-existing segments in polyps remains unknown.

TOR signaling is a major regulator of growth and RPS6 is an evolutionary conserved target of this pathway in eukaryotes^35,37^. Our findings show that tentacle development in polyps is preceded by a feeding-dependent global growth phase, followed by the formation of localized cell proliferation zones that define tentacle bud sites. Both patterns of growth correlate with the phosphorylation of RPS6, which is highly enriched in developing tentacles compared to the rest of the body. However, our results also suggest that distinct upstream inputs co-regulate cell proliferation and the phosphorylation of RPS6. These findings are consistent with the distinct sensitivity of these two processes to starvation. While the uniform pattern of cell proliferation is primarily dependent on feeding, the localized cell proliferation and phosphorylation of RPS6 at polyp tentacle primordia selectively requires *Fgfrb* function. We propose that *Nematostella* translates nutrient inputs into organismal growth, which in turn modulates the pre-defined developmental signalling landscape of body segments and promotes post-embryonic tentacle development. Interestingly, discrete ring muscle cells expressing *Fgfrb* pre-mark the sites of post-embryonic primordia and nucleate the early morphogenetic events associated with the feeding-dependent tentacle development. In the flatworm, muscle fibers can encode positional information that is critical for guiding tissue growth and regeneration^45,46^. While this property of muscles has not been reported in cnidarians, the feeding-dependent tentacle development in *Nematostella* offers the opportunity to explore the developmental function of ring muscles.

In contrast to tentacle development in polyps, cell proliferation is not spatially patterned during embryonic tentacle morphogenesis^4^. This difference suggests that there are distinct morphogenetic trajectories leading to embryonic and nutrient-dependent post-embryonic tentacle development. The *Nematostella* genome contains 2 FGF receptors (a and b) and 15 putative FGF ligands^38,40^. The function of the *Nematostella* FGF signaling pathway has only been investigated during embryonic and larval development^40^. Based on knockdown experiments, *Fgfra* is essential for apical organ formation and metamorphosis. In the current study, we generated a stable mutant line for *Fgfrb*. Phenotypic analysis of *Fgfrb* shows that mutant larvae undergo metamorphosis, but the process of axial elongation is disrupted. Interestingly, FGF signaling plays a critical role in the elongation of vertebrate embryos^47^. While further investigation is required to define the specific role of *Fgfrb* during embryonic *Nematostella* development, this finding highlights a potential pre-bilaterian function of FGFR signaling in body elongation. These results also reveal that *Nematostella* FGF receptors have distinct functions during development, suggesting the subfunctionalization of paralogs, with *Fgfra* having a dominant role during embryonic development. On the other hand, *Fgfrb* mutant polyps are viable and exhibit the four primary tentacles. With careful feeding and attention, these mutants can grow to some extent, but they do not show the localized cell proliferation and phosphorylation of RPS6 that are characteristic of polyp tentacle primordia. In sum, this finding reveals that FGF signaling couples the nutrient-dependent organismal growth with post-embryonic tentacle development.

## Materials and Methods

### Animal husbandry and stereomicroscopic imaging

*Nematostella vectensis* were cultured in 1/3 artificial seawater (Sea Salt; Instant Ocean). Adult polyps were spawned using an established protocol^15^. Progeny from nine spawning events were segregated into three groups, each containing approximately 1500 animals. Those groups were then further subdivided into more manageable populations of 250-350 animals. Each dish was fed *Artemia* nauplii 2-3 times per week. To ensure culture quality, seawater and dishes were changed weekly and biweekly, respectively. Tentacle pattern was monitored at different post-feeding time points and polyps were selected for fixation when new tentacle stages were observed. Selected polyps were relaxed in 7% MgCl_2_ (Sigma-Aldrich) and fixed in 4% paraformaldehyde (Electron Microscopy Sciences) for 90 minutes at room temperature. Larger animals were fixed in 4% paraformaldehyde overnight at 4°C. The fixed animals were washed with PBS and stored at 4°C until imaged. To orient the oral pole of fixed polyps along the directive axis, two morphological features were used. The position of primary mesenteries served as a landmark to orient polyps progressing from 4 to 12 tentacles. However, as the animals grow, this morphological feature became less pronounced in adults bearing more then 12 tentacles. At these stages, the location of siphonoglyph was more visible and was used to define the polarity of the directive axis. For imaging, polyps were decapitated and directly imaged in the Petri dish using a Leica MZ16F stereomicroscope with a QImaging QICAM *FAST* 12-bit color camera.

### Feeding of primary polyps and drug treatment

Prior to feeding, developing animals were raised for 3-4 weeks at 23°C in 1/3 artificial seawater in the dark. To perform feeding, ~300μl of concentrated *Artemia* was partially homogenized and mixed with 40-50 primary polyps in a 6cm Petri dish. One day of feeding corresponded to the incubation of polyps with *Artemia* for ~3 hours followed by a water change. For drug treatments, Rapamycin (1 μM; Sigma, R8781) and SU5402 (20 μM; Sigma, SML0443) were applied in 0.2% DMSO in artificial seawater at room temperature in the dark and were refreshed daily post-feeding. Concurrently, control animals were incubated in 0.2% DMSO in artificial seawater.

### Immunohistochemistry, EdU incorporation and lipid droplet staining

Polyps were incubated with EdU (300 μM from a stock dissolved in DMSO) in artificial seawater for 30 minutes (Click-it Alexa Fluor 488 Kit; Molecular Probes) as previously reported^48^. After incorporation, animals were relaxed in 7% MgCl_2_ in artificial 1/3 seawater for 10 minutes, fixed in cold 4% paraformaldehyde (Electron Microscopy Sciences) in 1/3 artificial seawater for 1 hour at room temperature. Samples were washed three times in PBS and permeabilized in PBT (PBS with 0.5% Triton X-100; Sigma) for 20 minutes. The reaction cocktail was prepared based on the Click-it Kit protocol and incubated with the animals for 30 minutes. After three washes in PBS, samples were labeled with Hoechst 34580 (1 μg/ml; Molecular Probes) in PBT overnight at 4°C. When combined with immunostaining, EdU-labelled polyps were stained with primary (rabbit anti-pRPS6 Ser235/236, 1:50; Cell Signaling #4858) and secondary (goat anti-rabbit IgG Alexa Fluor 594, 1:500; Molecular Probes) antibodies as previously described^4^. For phalloidin and DAPI staining, polyps were fixed in 4% paraformaldehyde with 10 mM EDTA for 1 hour and staining was carried out as previously reported^4^. For BODIPY staining, fixed animals were incubated with BODIPY^®^ 493/503 (1 μg/ml; ThermoFisher Scientific) for 60 minutes and then washed four times with PBS. To image the oral view of polyps, specimens were incubated in 87% glycerol (Sigma), decapitated with a sharp tungsten needle and mounted on glass slides with spacers. All images were taken with Leica SP5 or SP8 confocal microscopes. Oil Red O staining was performed as previously reported^49^ and imaged using LEICA MZ 16F microscope.

### RNA *situ* hybridization

The RNA *in situ* probe for *Fgfrb* was designed to cover ~ 900 nucleotides. cDNA of *Fgfrb* was cloned from total RNA isolated from mixed stages of animals using the RNeasy Mini Kit (Qiagen). Amplification primers were: *Fgfrb_fwd* 5’-AAACGCGAAAAGACCCTGATAGC-3’ and *Fgfrb*_rev 5’-GGACAGCGGGGACGTCAG-3’ Antisense probe was synthesized by *in vitro* transcription (MEGAScript Kit; Ambion) driven by T7 RNA polymerase with DIG incorporation (Roche). Chromogenic and fluorescent RNA *in situ* hybridization were carried out as previously described^50,51^. Fluorescent *in situ* hybridization coupled with immunostaining was performed as previously reported^52^. Bright field images were acquired under a Leica DM4000 microscope and confocal images were taken with a Leica SP8 microscope.

### CRISPR/Cas9 mutagenesis, transgenesis and crosses

CRISPR/Cas9 genome editing in *Nematostella* embryos was carried out as previously described^25^. Guide RNAs (gRNAs) were designed using the online web interface http://chopchop.cbu.uib.no. Two gRNAs targeting the first and second coding exons were generated via PCR reaction and purified using QIAquick PCR Purification Kit (QIAGEN). The sequence targets were: 1^st^ coding exon: GGTGCACGCCGTTTCACTGG**TGG** and 2^nd^ coding exon: **CCA**ATGTTTCGTGGCGCGCCCGC. gRNAs were *in vitro* transcribed using the MEGAshortscript T7 kit (Life Technologies) and purified using 3M sodium acetate/ethanol precipitation. Recombinant Cas9 protein with NLS sequence (800 ng/μl; PNA Bio, #CP01-20) was co-injected with each gRNA (500ng/μl) into unfertilized *Nematostella* oocytes. Injected oocytes were then fertilized and raised at room temperature.

To test the efficiency of genome editing in F0 injected-animals, genomic DNA was extracted from individual primary polyps following a previously described method^25^. We used 2μl of genomic DNA extract for subsequent PCR analysis. Following sequencing confirmation of indel or deletion events, F0 injected animals with tentacle defects were raised to sexual maturity and crossed to wild type animals. The progeny of these crosses was raised and individually genotyped using genomic DNA extracted from ~5 surgically isolated tentacles. Sequenced PCR products showing overlapping peaks in their chromatograms were cloned using NEB PCR cloning kit (NEB #E1202S) and then sequenced to define the identity of mutations carried by F1 heterozygous animals. For each gRNA-induced lesion, F1 animals carrying identical frameshift mutations were grouped by sex to form spawning groups. To confirm the relationship between genotype and phenotype, F2 progeny resulting from the crosses of F1 heterozygous animals were individually genotyped as described above.

The *Fgfrb-eGFP* transgenic line was generated by meganuclease-mediated transgenesis^26^. The genomic region surrounding the *Fgfrb* promoter (coordinates: Scaffold 4; 2003298-2012061) was cloned in the transgenesis plasmid^26^ and the first coding exon of *Fgfrb* was replaced by eGFP with the SV40 poly A using NEBuilder HiFi DNA assembly (NEB, E2621). F2 *Fgfrb-eGFP* homozygotes were established and crossed to *Fgfrb* heterozygous animals to combine the mutant alleles with the *Fgfrb* reporter construct. In all experiments, the expression of eGFP was visualized using immunostaining with Anti-eGFP antibody (mouse, ThermoFisher, A-11120).

### Quantification of EdU intensity and body dimensions

Images were analysed with ImageJ software^53^. EdU and Hoechst signals were quantified by the total intensity in a selected area of the oral pole spanning s3/s7 segments. The oral area was delineated with the straight line tool with 25μm width. For body dimensions, body length was measured from the oral opening to the aboral end. Body width was measured at the base of the tentacles. A straight line tool was used for both measurements. Oral circumference was defined with the oval selection in z-projection images of oral views.

## Acknowledgments

We thank the Stowers institute and EMBL aquatic cores for animal husbandry, as well as ALMF at EMBL for imaging support. We also thank A. Ephrussi, A. Aulehla and T. Hiiragi for discussion and comments on the manuscript. This work was supported by the Stowers Institute for Medical Research and European Molecular Biology Laboratory. Marie Anzo is supported by Human Frontier Science Program LTF (LT00126/2019-L).

## Author contributions

A.I designed and conceptualized the study, and wrote the manuscript. M.G. also designed the study and edited the manuscript. A.I generated the mutant and reporter lines. M.R.M. and A.I built the spatio-temporal map of tentacle addition. P.S. performed *in situ* hybridization experiments, genotyping and genetic crosses. P.S. and M.A. performed all drug-treatment and staining experiments. L.R.E. raised founders and heterozygous mutant animals to sexual maturity. A.S. performed image processing.

## Data Availability

We declare that the main data supporting the findings of this study are available within the article and its Supplementary Information files. Source data are available in the Source Data file for Figures 2–7, and Figures S2, S3, S5, S11 and S12. Extra data are available from the corresponding author upon request.

