## Supplemental information for "Feeding-dependent tentacle development in the sea anemone *Nematostella vectensis*"

Figure S1-S12

Supplemental movie 1 and 2

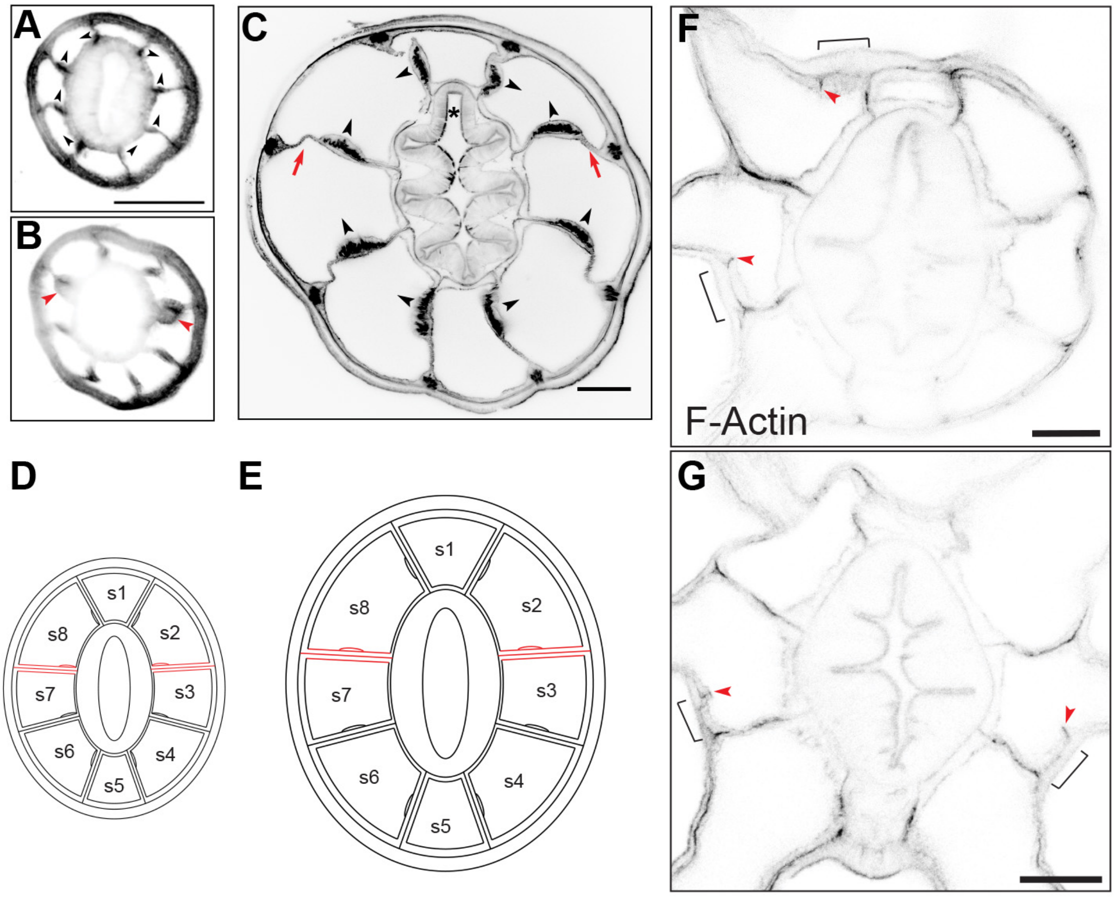


**Figure S1**: Body segments in primary and adult polyps. **A**, Cross-section through the oral pole at the base of the tentacles in a primary polyp. **B**, Cross-section through the oral pole of the same animal at the base of the pharynx and showing the most apical domain of primary mesenteries (red arrowheads). **C**, Cross-section through the oral pole of an adult polyp. Red arrows indicate the two primary mesenteries. Black asterisk (*) indicates the position of the siphonoglyph. All animals are labelled with Phalloidin (F-actin, in grey). Each mesentery bears a retractor muscle. The orientation of these muscles (black arrowheads) can serve as readout for the polarity of the directive axis. **D** and **E**, Diagrammatic cross-section through the oral pole showing the eight body segments in primary and adult polyps, respectively. Primary mesenteries are colored in red. Scale bares are 100µm in **A** and **C**. **F** and **G**, Confocal cross-sections of representative oral poles bearing eight tentacles and stained to label F-actin. The formation of short gastrodermis folds within segments precedes the third and fourth trans-budding events. Red arrowheads indicate new endodermal folds enriched in actin. Brackets show new tentacle territories. Scale bars are 50µm.

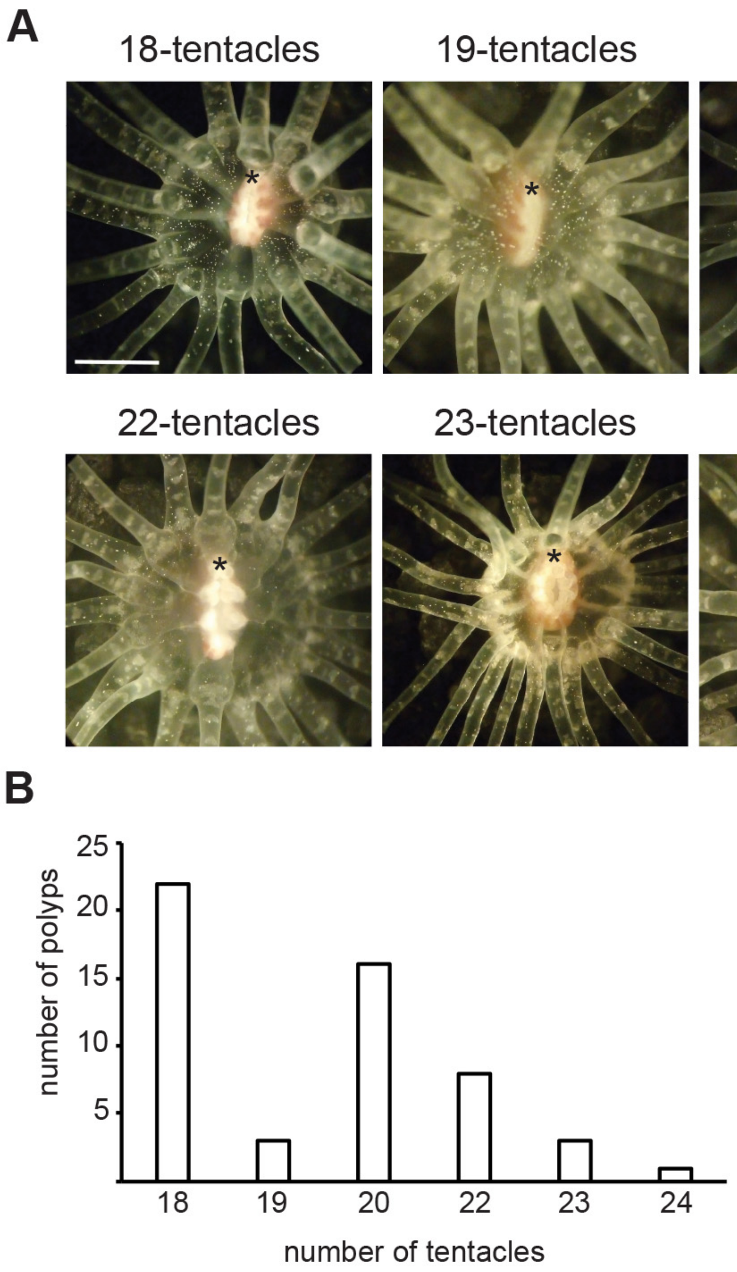


**Figure S2**: Adult polyps bearing more than 16-tentacles. **A**, oral views of adult polyps. Black asterisk (*) indicates the position of the siphonoglyph. Scale bar is 1mm. **B**, Quantification of polyps with indicated tentacle number. Source data are provided as a Source Data file.

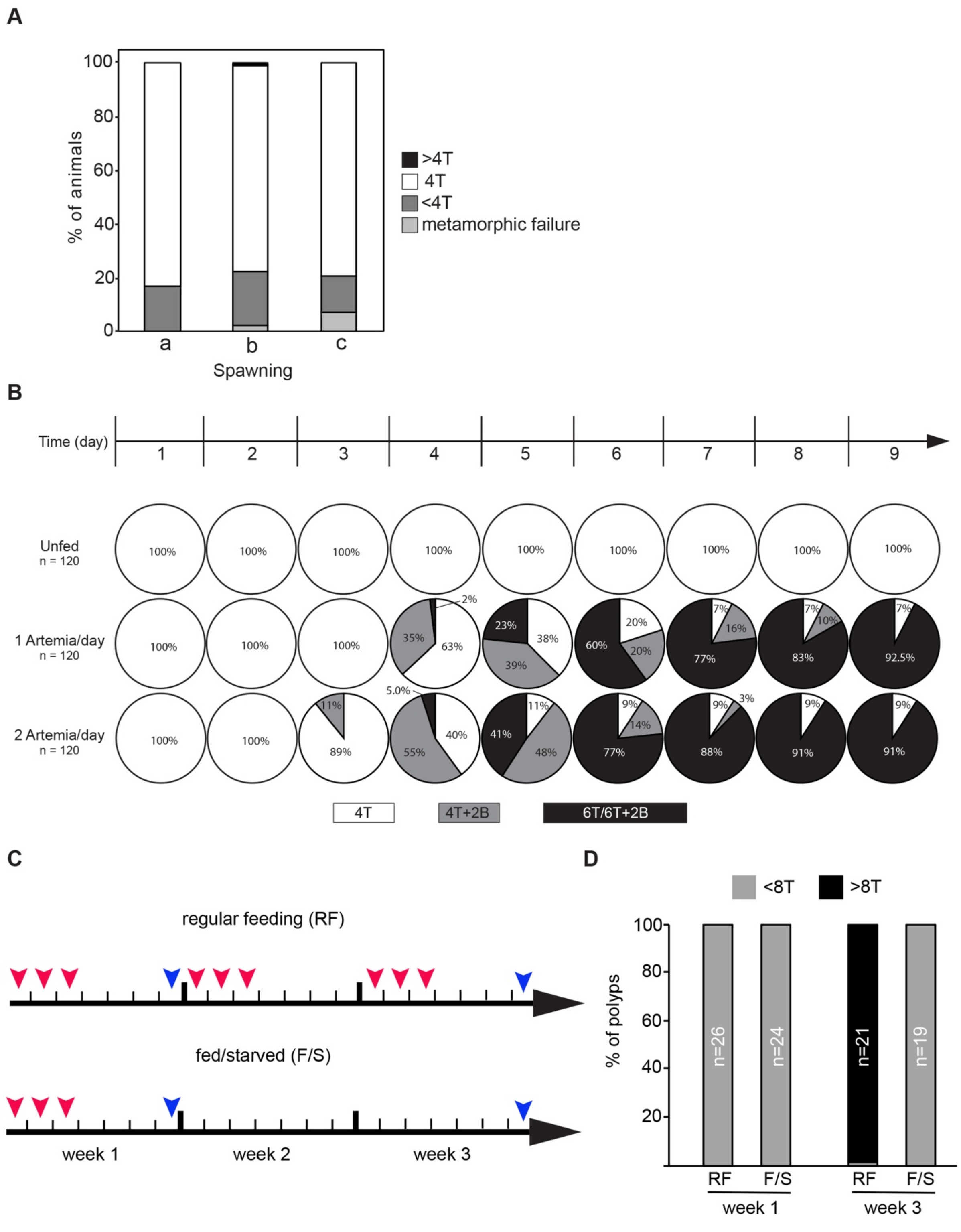


**Figure S3**: Feeding-dependent tentacle addition. **A**, Tentacle number in unfed primary polyps from three independent spawning: group a, n=101; group b, n=105; group c, n=100. Note that the frequent number of tentacles in unfed primary polyps is four. 4T: 4 tentacles; >4T: more than 4 tentacles; <4T: less then 4 tentacles. **B**, Feeding is required for subsequent tentacle addition in polyp stage. Unfed primary polyps did not add tentacles.

Polyps were hand-fed with 1 or 2 *Artemia* per day. Polyps fed twice developed buds earlier than those fed once per day. **C**, Tentacle addition in polyps that were fed for three weeks (regular feeding, RF) versus those that were fed for one week and then starved for 2 weeks (fed/starved, R/S). Red arrowheads indicate the feeding days. Blue arrowheads show the days where the tentacles were counted. **D**, Quantification of tentacle number in polyps exposed to these feeding treatments. Source data are provided as a Source Data file.

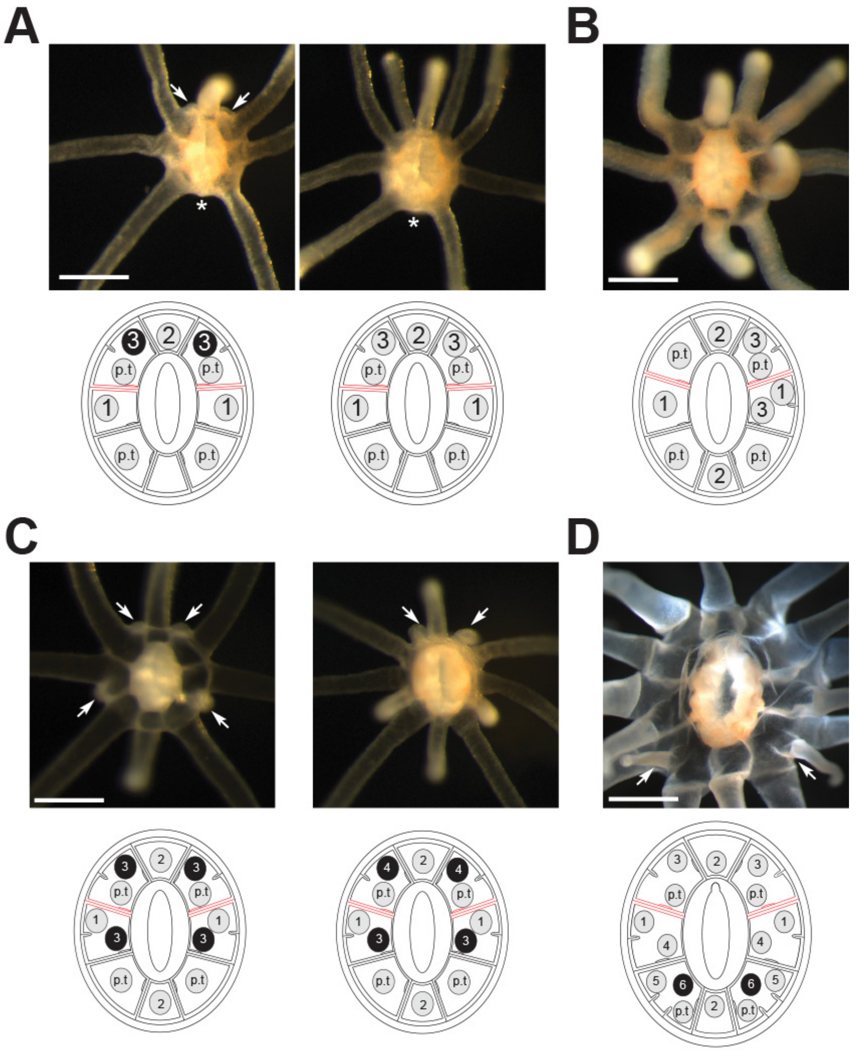


**Figure S4**: Variability in tentacle addition. **A**, One tentacle from the second *trans*-budding event was skipped while the polyp progressed to the third *trans*-budding. **B**, Tentacle development took place in neighbouring segments. **C**, Two pairs of buds formed simultaneously instead of sequentially. Scale bars are 250μm. **D**, A polyp showing a trans-budding event during phase III of tentacle addition. Scale bar is 500μm. White asterisk indicates a missing tentacle and white arrows show tentacle buds.

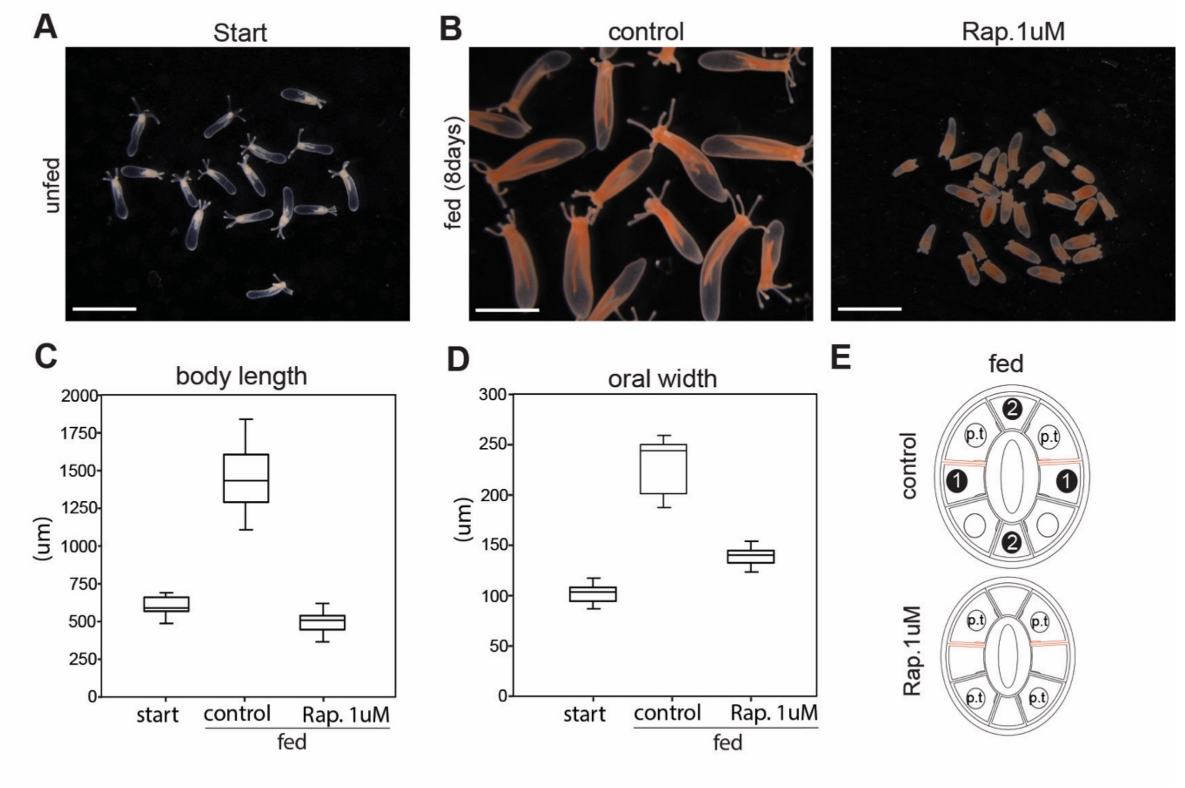


**Figure S5: A**, Unfed primary polyps. **B**, Control and Rapamycin (Rap.) treated polyps that were fed for 8 days. Note the orange color of fed polyps compared to unfed. Scale bars are 1mm. **C** and **D**, Quantification of body length and oral width of indicated conditions (start, n=13; control n=15; Rap. 1$\mu$M n=15). **E**, Diagrams showing tentacle arrangement in fed controls and Rapamycin treated animals. Source data are provided as a Source Data file.

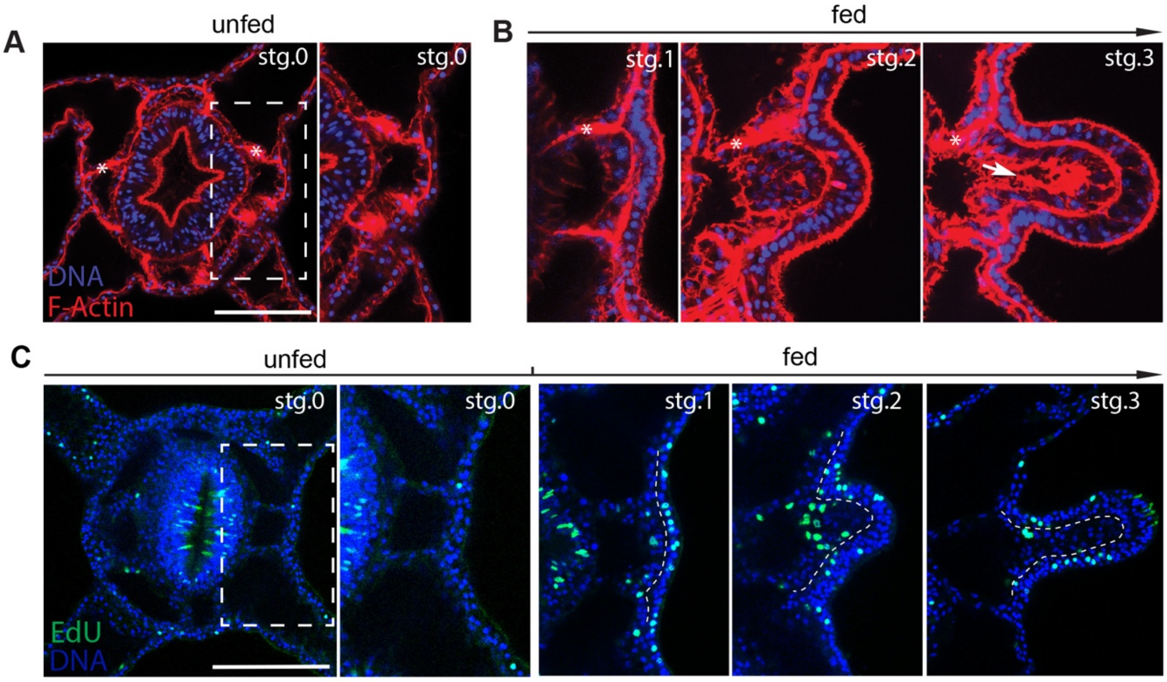


**Figure S6: A,** Confocal z-section of the oral pole of an unfed polyp stained with phalloidin to visualize F-Actin (red) and Hoechst to label nuclei (blue). Inset box is a zoom-in of s3 segment. **B**, Confocal sections of developing buds at sequential stages. Budding initiates from a flat epithelial architecture (stage 0; stg.0) that undergoes a slight asymmetric thickening towards the primary mesentery (stage 1; stg.1) followed by the formation of an outgrowth (stage 2; stg.2). A mature bud (stage 3; stg.3) is formed when a lumen appears (white arrow). **C**, Cross-sections of animals stained for EdU incorporation (green) and with Hoechst (blue) to visualize S-phase cells and nuclei at the indicated feeding conditions. Inset box is a zoom-in of s3 segment. Dashed lines separate the outer- and inner-layer nuclei. Scale bar is 50μm.

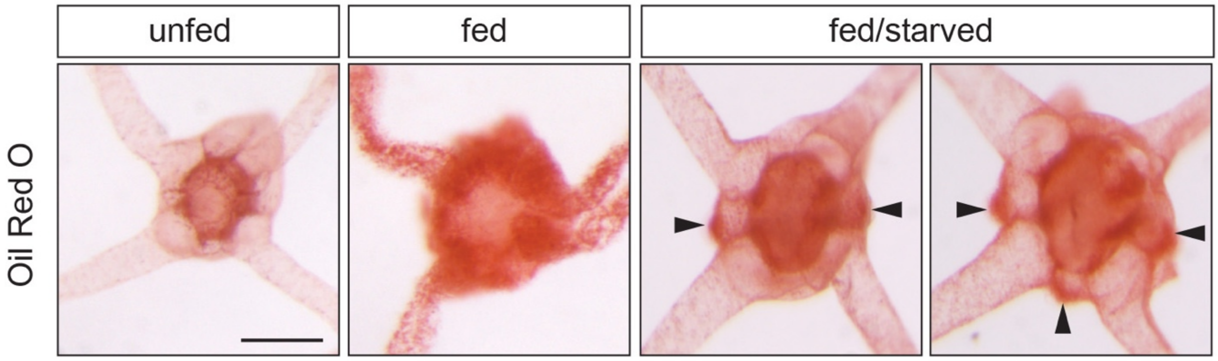


**Figure S7**: Oral views of animals stained with Oil Red O at the indicated feeding conditions. Arrowheads indicate the enrichment of Oil Red O staining in tentacle primordia. Scale bar is 50μm.

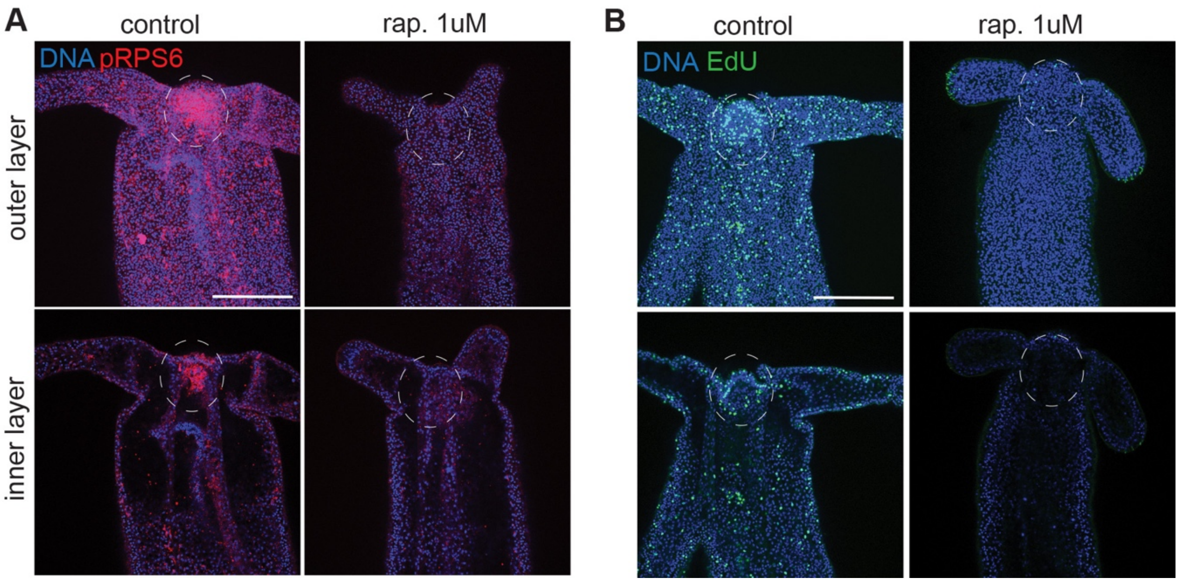


**Figure S8**: **A**, Confocal projections of lateral views of 4 days fed control and Rapamycin-treated polyps stained with an antibody against pRPS6 (red) and Hoechst to label nuclei (blue). Rapamycin was added at day 3 and incubated for 24 hours. **B**, Confocal projections of lateral views of 4 days fed control and Rapamycin-treated polyps stained for EdU incorporation (green) and Hoechst to label nuclei (blue). Top image is surface view and bottom image is a cross-section. Dashed circles indicate the location of tentacle primordia in the outer and inner layers. Scale bars are 50μm.

```
Human           MKLNISFPATGCQKLIEVDDERKLRTFYEKRMATEVAADALGEEWKGYVVRISGGNDKQG 60
Nematostella    MKLNISYPVTGCQKLIEIEDERKVRSFYDKRMAMEVSGECLGDEWKGYVFRITGGNDKQG 60
Yeast           MKLNISYPVNGTQKTIEVDDEHRVRVFYDKRIGQEVNGEAVGDEFKGYVFKIAGGNDKQG 60
                ******:*..* ** **::**:::* **:**:. ** .:.:*:*:****.:*:*******

Human           FPMKQGVLTHGRVRLLLSKGHSCYRPRRTGERKRKSVRGCIVDANLSVLNLVIVKKGEKD 120
Nematostella    FPMKQGIMTNGRVRLLLSKGHSCYRPRRTGERKRKSVRGCIVDSQLSVLSLVIVKKGEQD 120
Yeast           FPMKQGVLLPTRVKLLMAKGTSCYRPRRNGERKRKSVRGAIVGPDLAVLALIITKKGDQE 120
                ******::   **:**::** *******.**********.**..:*:** *:*.***:::

Human           IPGLTDTTVPRRLGPKRASRIRKLFNLSKEDDVRQYVVRKPLN-KEGKKPRTKAPKIQRL 179
Nematostella    IPGLTDNTIPRRLGPKRVGKIRKMFNLSKEDDVRQYVIRRPLPEKEGKKAKSKAPKIQRL 180
Yeast           IEGITNESVPKRLGPKRANNIRKFFGLTKDDDVRDFVIRREVV--KGDKTYTKAPKIQRL 178
                * *:*: ::*:******...***:*.*:*:****::*:*: :   :*.*. :********

Human           VTPRVLQHKRRRIALKKQRTKKNKEEAAEYAKLLAKRMKEAKEKRQEQIAKRRRLSSLRA 239
Nematostella    VTPVVLQRKRKRLALKRQRAQKCKQEAADYAKLLAKRAKEAKEKRHEQLMKKRRASSLRD 240
Yeast           VTPQRLQRKRHQRALKVRNAQAQREAAAEYAQLLAKRLTEKKAEKAE--ERKRRASSLKA 236
                ***  **:**:: *** :.::  :: **:**:***** .* * :: *   ::** ***:

Human           STSKSESSQK 249
Nematostella    SVSAK----- 245
Yeast           ----------
```

**Figure S9:** Protein sequence alignment of ribosomal protein S6 from human, *Nematostella* and yeast. Asterisks (*) show conserved residues. Colons and periods indicate conservation of groups of strongly and weakly similar properties, respectively. The phosphorylated serines in human are conserved and highlighted in green.

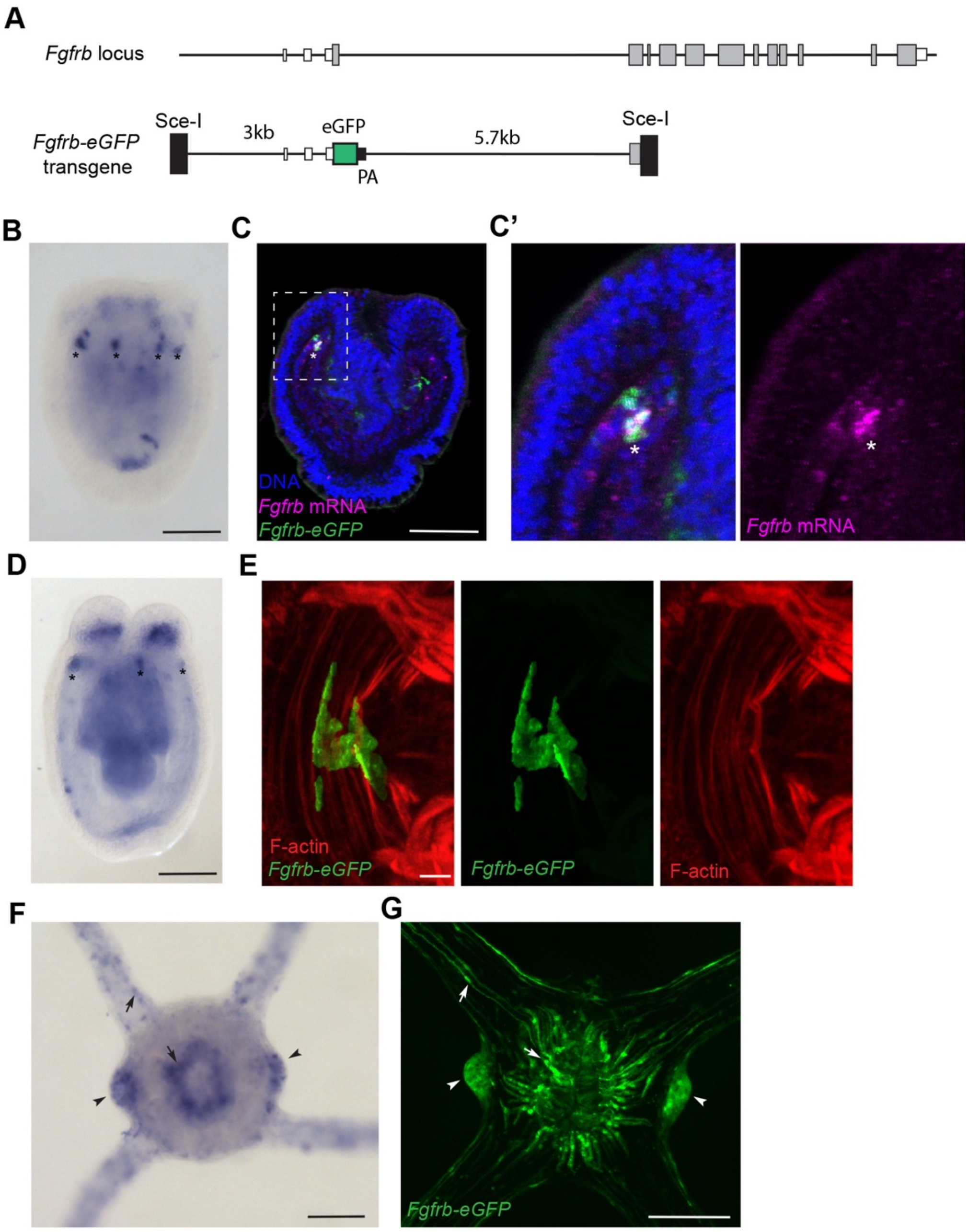


**Figure S10**: **A**, Gene model of the *Fgfrb* locus and map of the *Fgfrb-eGFP* transgene. **B**, **D** and **F**, *In situ* hybridization of *Fgfrb* in developing *Nematostella*. **B**, Planula. **D**, Unfed polyp. **F**, Oral views of a fed polyp. Asterisks indicate clusters of *Fgfrb*-expressing cells in the gatrodermis. **C,** Confocal z-projections of the *Fgfrb-eGFP* transgenic planula stained with $\alpha$-eGFP (green) and Hoechst (blue) as well as labeled for *Fgfrb* mRNA (purple). **C'**, Inset box is a zoom in view of panel C. **E**, Confocal z-projections of the *Fgfrb-eGFP* transgenic polyp stained with $\alpha$-eGFP (green) and Phalloidin (red) showing a high magnification of an *Fgfrb*-positive cell. **G**, Confocal z-projections of a budded *Fgfrb-eGFP* polyp. Arrowheads indicate tentacle buds. Arrows shows longitudinal muscle cells. Scale bars are 50$\mu$m in B, C, D, F and G. Scale bar is 3$\mu$m in E.

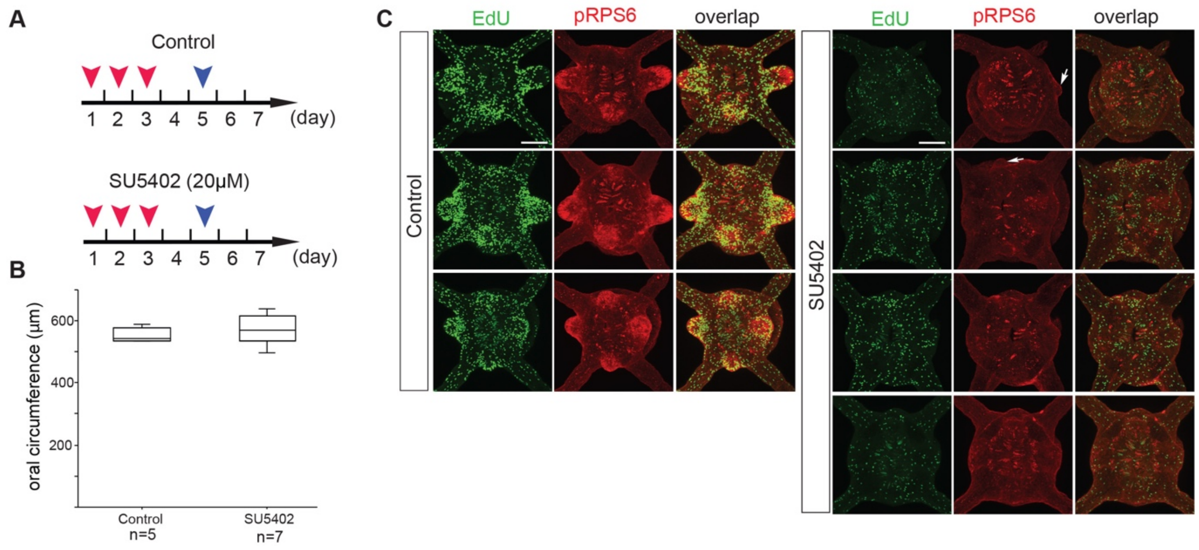


**Figure S11**: **A**, Feeding assay for controls and SU5402-treated polyps. Red arrowheads indicate the feeding days. Blue arrowheads show the days where animals were fixed. **B,** Quantification of oral circumferences in the indicated conditions. Box plot values consist of the median (center line), upper and lower quartiles (upper and lower edges of box), and maximum and minimum values (whiskers). Outliers are represented in small circles. **C,** Confocal z-projections of oral poles of indicated polyps stained for EdU incorporation (green), an antibody against pRPS6. Arrows indicate reduced outgrowth forming in few SU5402-treated polyps. Scale bars are 50μm. Source data are provided as a Source Data file.

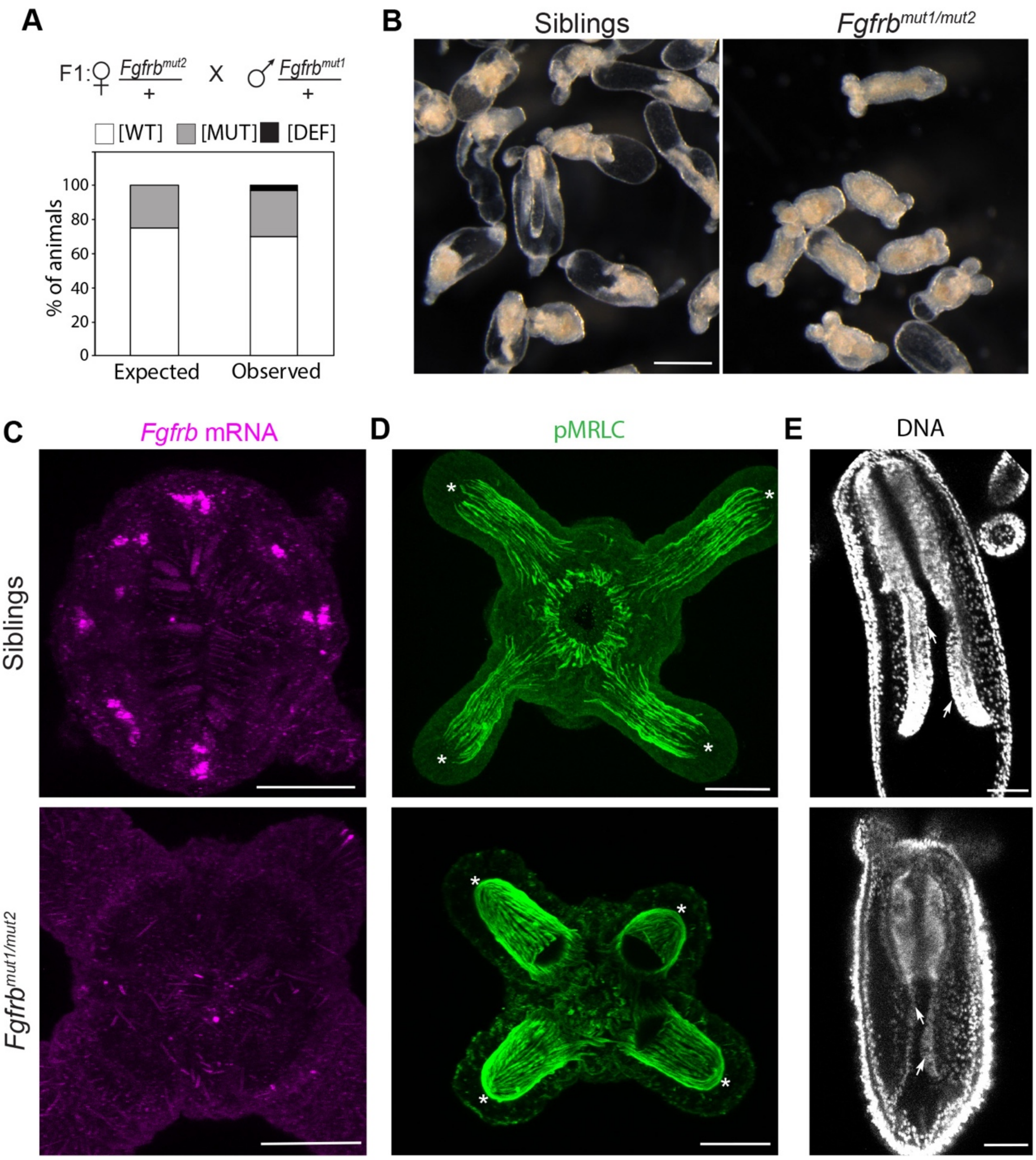


**Figure S12**: **A**, F1 heterozygotes cross and quantification of F2 polyp phenotypes (n=156). These include wild type (WT), mutant (MUT) and deformity (DEF) phenotypes. **B**, Images of *Fgfrb*$^{mut1/mut2}$ polyps and their siblings. Scale bar is 250μm. **C**, Fluorescent *in situ* hybridization of *Fgfrb* (purple) in a wild type and the *Fgfrb* mutant polyp. **D**, Immunostatining of phospho-Myosin Regulatory Chain (pMRC, green) labelling tentacular and oral muscles in a wild type and the *Fgfrb* mutant polyp. Asterisks (*) indicate longitudinal muscle organization at the tentacle tips. Note that longitudinal muscles project through the tip region in the *Fgfrb* mutant while they do not cross in the wild type. **E**, Confocal z-projections of the body column of indicated polyps stained with Hoechst (white). White arrows show septal filaments. The *Fgfrb* mutant polyp has reduced septal filaments compared to the control. Scale bars are 50μm. Source data are provided as a Source Data file.

Movie 1: 3D view of an oral pole showing *Fgfrb* expression (green) visualized by mRNA *in situ* hybridization and trained nuclei (blue). The 3D view was reconstructed using Imaris x64 9.2.1 (full ref: Imaris v 9.2.1, ImarisXT, Bitplane AG, software available at http://bitplane.com)

Movie 2: 3D view of the oral pole for the *Fgfrb-eGFP* transgenic line stained with α-eGFP (green) and Hoechst (blue).
